# Antibiotics Rewire Core Metabolic and Ribosomal Programs in Mammalian Cells

**DOI:** 10.1101/2025.06.20.660816

**Authors:** Cameron S. Movassaghi, Jesse G. Meyer

## Abstract

Antibiotics are routinely added to mammalian cell culture media to prevent bacterial growth. However, the use of antibiotics in cell culture can confound downstream experimental results. While genomic and transcriptomic differences between cell cultures treated with and without antibiotics are well-documented, far fewer, if any, comprehensive proteomic comparisons on the use of antibiotics in cell culture have been performed. Here, we present a study on the proteome-wide differences of culturing HepG2 cells in antibiotic (*i.e.,* penicillin/streptomycin) and non-antibiotic-containing media. Using a longitudinal and crossover treatment study design, we analyzed 119 samples across nine passages and four conditions. On average, 9,374 proteins were detected per sample, and we identified 383 proteins that were differentially abundant between conditions. These changes included ribosomal and mitochondrial proteins, demonstrating that off-target effects of antibiotics on mammalian cells occur at the protein level. Linear mixed-effects modeling suggested that the proteomic impact of antibiotic treatment is strongest in the first passage after treatment and stabilizes after approximately three passages. Furthermore, initiating antibiotic treatment induced a greater number of differentially abundant proteins than discontinuing treatment. Lastly, we compared our results to existing literature on the use of common antibiotics in mammalian cell culture. We identified proteins and pathways conserved across studies, -omics layers, and cell types. We hope that this detailed proteomic survey will aid researchers in comparing cross-study or cross-condition results from antibiotic-treated mammalian cells and inform appropriate experimental designs for the use of antibiotics in cell culture.

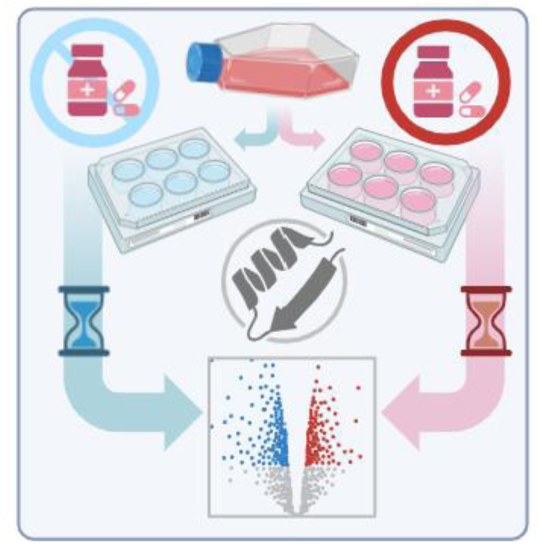

## INTRODUCTION

Mammalian cell culture media is commonly supplemented with antibiotics for their bacteriostatic or bactericidal effects.^1^ Regardless of an active bacterial contamination, antibiotics (*e.g.,* penicillin/streptomycin; PenStrep) are often added by default to lower the risk of future contamination. Although chronic treatment of cell cultures with antibiotics is generally accepted practice, others have advocated instead for strict aseptic techniques and the use of antibiotics only when contamination is suspected.^2–4^ Chronic reliance on antimicrobials may lead to poor adherence to aseptic protocols, allowing latent contamination to go unnoticed and increasing the risk of drug-resistant microbes.

Studies on the unintentional effects of antibiotic supplementation on cell properties have emerged as an alternative reason to forgo such treatment.^3–9^ Antibiotics exposure is linked to changes in cell growth, proliferation, differentiation, as well as cytotoxic and apoptosis-inducing properties.^4,9^ At high concentrations, they can inhibit protein synthesis and affect enzymatic activity as early as four hours after treatment.^5,7^ At the genomic level, Ryu *et al.* used RNAseq to identify over 200 differentially expressed genes in HepG2 cells grown with *versus* without PenStrep.^3^

While this evidence suggests the proteome would also be affected, few studies have examined the impact of antibiotic treatment of cell cultures at the protein level. Of these studies, the panels of measured proteins ranged from dozens to hundreds,^6^ which is relatively small in comparison to recent proteomic methods capable of identifying over 10,000 proteins.^10^ Given the ubiquity with which cell culture media is supplemented with antibiotics for *in vitro* proteomic experiments, a broader investigation into the proteome-wide effects is warranted. Antibiotics may dysregulate the proteins or pathways of interest in an *in vitro* model, leading to follow-up or translational studies that are confounded by the extent and type of antibiotic treatment. Conflicting studies also hinder integrative approaches to multi-omics, as data from varying antibiotic treatments are not considered. In summary, while the choice of antibiotic supplementation in cell culture is ostensibly a simple experimental variable, the downstream effects on biologically relevant changes to the proteome remain largely unexplored.

Pen/Strep is one of the most common antibiotic treatments for chronic supplementation in mammalian cell cultures. Their mode of action and effects on bacteria are well understood: penicillin is a β-lactam that inhibits cell-wall synthesis, while streptomycin is an aminoglycoside that targets bacterial ribosomal subunits.^2^ They are often used in combination due to their synergistic properties, *e.g.,* inhibition of cell wall synthesis by penicillin may allow easier access of streptomycin to bacterial ribosomes.^11^ However, both penicillin and streptomycin have documented toxic and other off-target effects in eukaryotic cells, the exact mechanisms of which are still being elucidated.^12^ For example, aminoglycosides have been recently shown to bind to human rRNA^13^ and as many as 75% of mammalian mitochondrial proteins are of bacterial evolutionary origin.^14^ Thus, antibiotics likely perturb mitochondrial and ribosomal proteins in eukaryotes. As such, dysregulation of protein synthesis and metabolic pathways resulting from antibiotic treatment alone may be confounded by other biological variables.^8^

Here, we measured proteome remodeling induced by penicillin and streptomycin in HepG2 cells. HepG2 cells were chosen due to their ease of handling, rapid proliferation, and suitability for drug response studies.^15,16^ We employed a longitudinal study design with treatment crossover to ensure robust results. In total, we detected approximately 10,250 proteins from 119 samples across nine passages and four treatment groups. As many as 139 proteins with Benjamini-Hochberg adjusted p-values<0.01, or 383 proteins at adj. p-values<0.1 (to be consistent with transcriptomics from Ryu *et al.*), were differentially abundant between the four conditions. These results were compared in a survey of relevant literature, revealing several conserved proteins and pathways that are consistently affected across -omic layers, such as proteostasis, transmembrane transport, ribosome biogenesis, and lipid metabolism. Notably, we observed a discordance between transcriptomic and proteomic changes, suggesting that the effects of antibiotic exposure may be context-specific and modulated by factors such as passage number, cell type, dosage, and treatment history. We hope that these results and the corresponding dataset can provide the proteomics community with a deeper understanding of the systematic effects of antibiotic supplementation in cell culture and help guide researchers in accounting for these effects in their *in vitro* experimental designs.

## RESULTS & DISCUSSION

### Experimental design

The experimental overview is shown in **Fig. 1**. Briefly, two six-well plates of HepG2 cells (seeded at the same density and from the same passage) were grown in parallel in antibiotic (A) and non-antibiotic (N) media (*i.e.,* 1% PenStrep; see *Methods*). On each passage, at least one well was used to seed the following passage, while the remaining wells were washed, lysed, and frozen until the day of analysis. Throughout, cells were seeded at approximately equal densities (∼0.2 × 10^6 cells per well) and harvested at equal confluency (>80% or ∼1.2 × 10^6 cells), as confirmed visually using an inverted microscope. In the fifth passage of the parallel A and N treatments, each condition was further split into two subgroups (a continued treatment group or a reverse treatment group) for four additional passages. Continued treatment refers to the use of the same media conditions, *e.g.,* A_1-9_, N_1-9_. Reverse, or crossover, treatment refers to switching the media to the opposing treatment; for example, cells grown in antibiotics for five passages were then grown in non-antibiotic-containing media for four passages (A_5_N_1-4_). This experimental design enabled us to continue monitoring a single condition for an extended period (29 days; 9 passages: A_1-9_, N_1-9_), while also replicating treatment transitions in both directions (A_5_N_1-4_, N_5_A_1-4_). On the same experimental day, all lysates were then thawed and underwent solid-phase enhanced sample preparation (SP3).^17^ Samples were then frozen until analyzed by liquid chromatography-tandem mass spectrometry with data-independent acquisition (see *Methods*).

**Figure 1.**
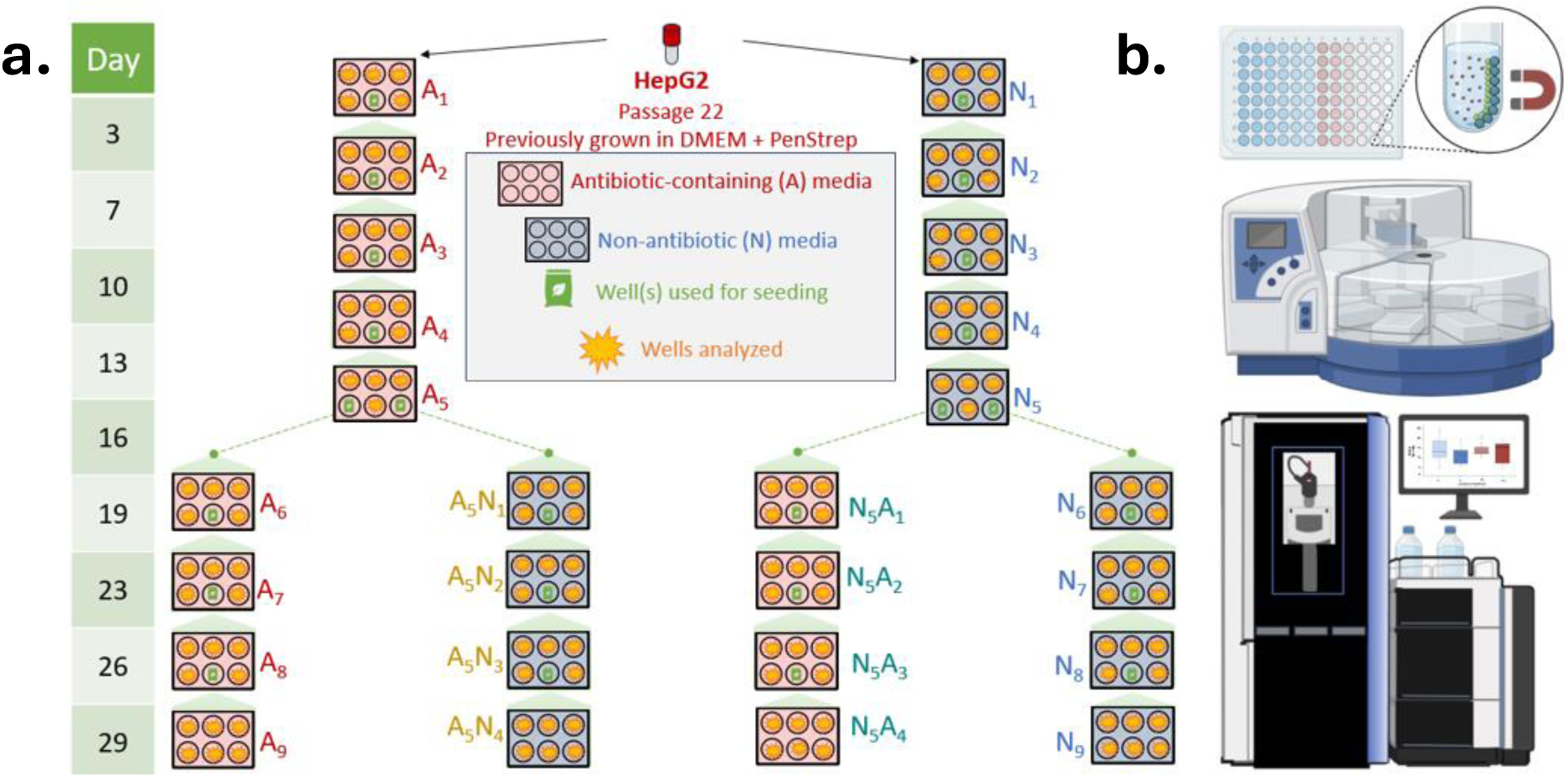
Experimental overview. **a)** HepG2 cells were grown in Dulbecco’s Modified Eagle Medium (DMEM) containing penicillin/streptomycin (PenStrep). At passage 22, a confluent T-75 flask was split into two six-well plates and grown in the same media, except that one plate contained antibiotics (A) and the other did not (N). Each plate was grown under the same conditions for five passages; the subscript represents the number of passages after splitting the cells into each treatment group *(i.e.,* A_1-5_, N_1-5_). After five passages, A_5_ and N_5_ were split again, each into two 6-well plates. The treatment either remained constant and grew for an additional four passages (i.e., A_6-9_, N_6-9_), or was swapped to the opposite condition (i.e., N_5_A_1-4_, A_5_N_1-4_). For each passage, one or two wells were used to seed the subsequent passage plate while the remaining wells were analyzed using the proteomics workflow. **b)** The proteomics workflow involved using single-pot, solid-phase enhanced sample preparation, followed by liquid chromatography-mass spectrometry with an Orbitrap Astral.

This study design has the following advantages. First, it enables the longitudinal tracking of proteomic changes across nine sequential passages, in contrast to prior studies that typically examined only single-passage time points or short-term antibiotic exposure (**Table S2**). By crossing the conditions (*i.e.,* crossover design), we increase the robustness and statistical power of our study by capturing both onset and withdrawal effects of antibiotic treatment within the same experimental framework. Changes in protein abundance that remain consistent across conditions and passages are likely to reflect generalizable biological responses, rather than transient effects due to passage-specific variation or genetic drift.^18^

### Global protein differences

The protein quantities were consistent across all samples (**Fig. 2a**). Approximately 9,200 proteins were detected on average (**Fig. 2b**). A total of 10,255 proteins were detected, and on average, each protein was detected in 108/119 samples (91%; **Fig. 2c**). Remarkably, 7,606 proteins were detected in all samples. The most abundant proteins included those related to cytoskeletal organization, stress response, and metabolism (**Fig. 2d**). Taken together, these results are expected, given the consistent handling and treatment of the cells, aside from the intended variation of antibiotic supplementation.

**Figure 2.**
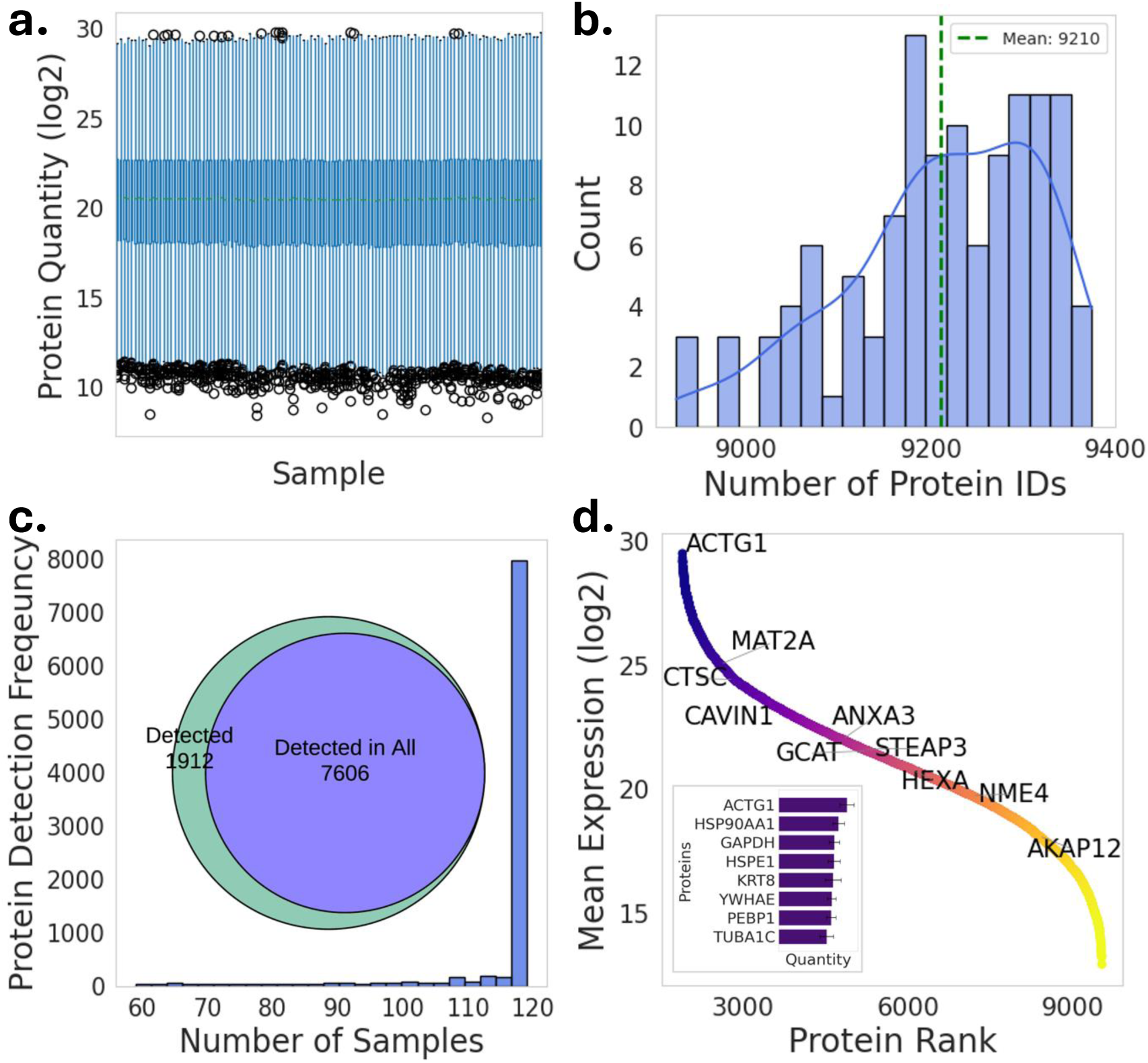
Protein quantification overview. **a)** Boxplot of protein quantities for all samples. **b**) Histogram of the number of detected proteins across all samples. **c)** Histogram of the number of samples in which each protein was detected. *Inset*: Euler diagram of all detected proteins *versus* the proteins detected in all samples. **d)** Average quantities of proteins across all samples. *Inset*: top eight most abundant proteins. All values are prior to imputation.

We then analyzed the variations in total protein quantities across plates, treatment conditions, and passages (**Figs. S1, S2 and Supplementary Note 1**). All variation was consistent across these experimental variables. A slight decrease in protein content was noted for specific wells of passages 5 and 6 due to variations in confluency, but this did not significantly alter any downstream results (see **Figs. S3-S5**).

### Linear mixed-effect modeling

Previous studies comparing antibiotic treatments on cultured cells have used *t*-tests or similar analyses to compare the log-fold changes of genes or proteins across treatment (A) *versus* control (N) at a single time point or passage.^3,8^ For direct comparison to these studies, we provide a discussion of *t*-test results in **Figs. S6-S9, Supplementary Note 2,** and **Table S1**. Interestingly, we found that moving from antibiotic-free cultures to antibiotic-containing cultures (N→A) consistently resulted in more differentially abundant proteins than moving from antibiotic-containing cultures to antibiotic-free cultures (A→N).

Given the longitudinal, multi-passage nature of our dataset, these pairwise time-point comparisons risk obscuring meaningful biological variation with passage-dependent variation. To address this, we utilized a linear mixed-effects model (LME). LMEs are useful for grouped datasets to incorporate both fixed effects (*i.e.,* parameters expected to vary systematically across an entire population) and random effects (*i.e.,* related to individualized experimental units or subjects).^19^ In our design, we considered the two primary sources of variation (treatment condition and passage) as fixed and random effects, respectively (**Fig. 3a**). Further discussion on the motivation and selection of LMEs compared to other linear models or statistical analyses is provided in the *Supplemental Information* (**Supplementary Note 3**).

**Figure 3.**
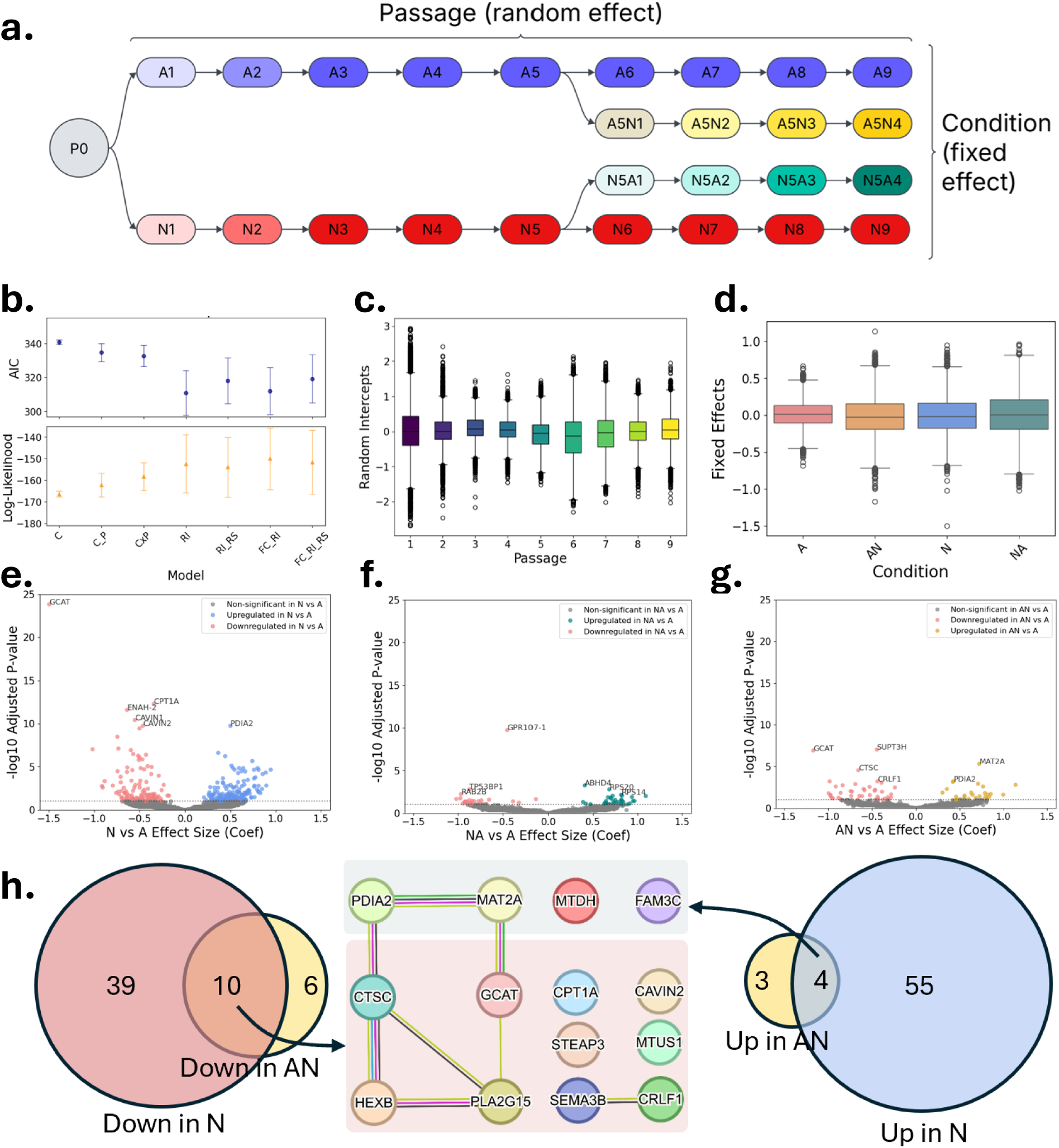
Linear mixed effect model reveals significant protein differences across conditions and passages. **a)** Flowchart distinguishing fixed *versus* random effects in the context of treatment condition *versus* passage. **b)** Model comparison metrics for model selection, where AIC represents Akaike information criterion. C, C_P, and CxP present ordinary least squares models with covariates of condition, condition + passage, and condition*passage, respectively. RI and RI_RS represent random intercept and random intercept + random slope linear mixed effect models, respectively, where passage is the random effect. FC_RI and FC_RI_RS are the same as the latter models, except condition has been added as a fixed effect. **c)** Boxplot of random intercepts for the FC_RI model. **d)** boxplot of the fixed effects for the FC_RI model. **e-g)** Volcano plots for each condition, where effect is given by the FC_RI fixed effect coefficient. An adjusted p-value using Benjamini-Hochberg correction was set to a p ≤ 0.01 cutoff. **h)** Euler diagrams for the proteins differentially abundant in each condition (left/right). STRING diagram for proteins consistently differentially abundant across conditions (middle).

### Differential expression

To determine the most appropriate linear model for our data, we evaluated multiple models using standard model selection criteria (**Fig. 3b**). The best-performing model was an LME with treatment condition as a fixed effect and passage as a random intercept (FC_RI; see **Supplementary Note 3** and **Figs. S10-S12**). The distributions of the random intercepts demonstrated that passage-by-passage variation was greatest in the first passage directly after a condition crossover (passages 1, 6), but stabilizes after roughly three passages (passages 3, 8) (**Fig. 3c**). These rapid changes in many individualized protein coefficients reflect transient proteome remodeling following changes to antibiotic exposure.

Using the FC_RI model, the fixed effect coefficients (**Fig. 3d**) can then be used to perform differential expression analysis of proteins across conditions (**Fig. 3e-g**). Here, we use A as the reference, as antibiotic-containing media is commonly used by default, as was the case in this study, and compare against N (no antibiotics), NA (crossover from N to A), and AN (crossover from A to N), respectively. In **Figure 3e**, positive coefficients are higher in N, and negative coefficients are higher in A. The y-axis represents adjusted p-values using the Benjamini-Hochberg procedure with a significance threshold of ≤ 0.01, unless otherwise noted.

Using this threshold, we found 108 differentially abundant in N *versus* A (59 up, 49 down), 8 proteins differentially abundant in NA *versus* A (5 up, 3 down), and 23 proteins in AN *versus* A (7 up, 16 down) (**Fig. 3h**). Across these three comparisons, this resulted in 139 unique differentially abundant proteins (71 up, 68 down, relative to A). Of these, FAM3C, MAT2A, MTDH, PDIA2 were increased in both N and AN (*i.e.,* consistently higher abundance when not exposed to antibiotics). Meanwhile, CAVIN2, CPT1A, CRLF1, CTSC, GCAT, HEXB, MTUS1, PLA2G15, SEMA3B and STEAP3 were decreased in both N and AN (*i.e.,* consistently higher abundance when exposed to antibiotics, **Fig. 3h**, center). The full list of all proteins differentially abundant across conditions is provided in **Supplemental Table 1**.

For brevity, we focus on the 14 select proteins differentially abundant across multiple comparisons, which is a benefit of our crossover design (**Fig. 3h**, center). The differentially abundant protein with the lowest adjusted p-value and greatest fixed effect coefficient was glycine C-acetyltransferase (GCAT). This protein was consistently more abundant in the presence of antibiotics. GCAT is a mitochondrial enzyme involved in generating glycine as part of the threonine degradation pathway. Glycine is one of the three amino acids required to synthesize the endogenous tripeptide glutathione, which plays a crucial role in the oxidative stress response, metabolic detoxification, and immune system regulation.^20^ Thus, the increase of GCAT in response to antibiotics may be implicated in oxidative stress protection mechanisms. Few studies have investigated GCAT, but some evidence suggests that this enzyme, along with others related to glycine metabolism, is implicated in metabolic regulation and aging.^21–23^

Along this same pathway related to one-carbon metabolism, methionine adenosyltransferase 2A (MAT2A) catalyzes the synthesis of S-adenosylmethionine (SAM) from methionine. SAM is required for the trans-sulfuration pathway to generate cysteine, which is then required for glutathione production. Glycine, and potentially GCAT activity, feed into the methionine cycle.^24^ However, unlike GCAT, MAT2A was found to decrease in the presence of antibiotics. While GCAT localizes to the mitochondria, MAT2A resides primarily in the nucleoplasm and cytoplasm, and is more active in extrahepatic tissues. Due to the production of SAM, MAT2A is implicated in methylation, which in turn affects gene expression and post-translational modifications. Its presence in the nucleus and cytoplasm could also indicate effects on ribosomal biogenesis and function. Regardless, these results suggest a regulatory effect of glycine and glutathione-related metabolism.

Changes in protein folding and lipid oxidation pathways parallel this shift in redox-related metabolism. Protein disulfide-isomerase A2 (PDIA2) was found to decrease in the presence of antibiotics. PDIA2 is located in the endoplasmic reticulum and is believed to function as a protein folding chaperone. Carnitine palmitoyltransferase 1A (CPT1A), found on the outer mitochondrial membrane, is a rate-limiting enzyme involved in fatty acid beta-oxidation. CPT1A was in higher abundance in the presence of antibiotics. While we hypothesized changes to mitochondrial proteins would occur, this specific alteration of a crucial lipid metabolism protein was unexpected.

Several membrane-associated proteins were also found to be differentially abundant. Caveolae-associated proteins (CAVINs), which contribute to membrane curvature and scaffolding, were decreased in response to antibiotics. These proteins have various roles associated with everything from the cytoskeleton to ribosomes, transcription, lipid metabolism, hypoxia, and cell signaling.^25^ Six-transmembrane epithelial antigen of prostate 3 (STEAP3) is another membrane protein that is also decreased in AN and N, with multiple roles in liver health and disease.^26^

Other robustly affected proteins include cytokine receptor-like factor 1 (CRLF1) and semaphorin-3B (SEMA3B), which are secreted proteins that regulate immune responses and apoptosis.^27,28^ The remaining proteins are associated with hepatocellular carcinoma, cell proliferation, and tumor growth, *including MTUS1 (microtubule-associated scaffold protein 1), metadherin (MTDH), and FAM3C (family with sequence similarity 3, metabolism-regulating signaling molecule C*). For example, MTUS1 is thought to be a tumor suppressor such that downregulation is associated with poor prognosis.^29^ Meanwhile, MTDH^30^ and FAM3C^31^ are known to be upregulated in various cancers, including hepatocellular carcinoma. In our data, MTUS1 was consistently increased in antibiotics while MTDH and FAM3C were consistently decreased with antibiotics. These and other differences reflect existing literature detailing the benefits and risks of antibiotics in cancer treatment and progression.^32^

### Pathway analysis

To better understand the biological pathways affected by antibiotic treatment, we performed gene ontology (GO) enrichment analyses on the differentially abundant proteins (**Fig. 4**). To assess general pathway trends, we grouped proteins that were more abundant in either A or NA, or less abundant in AN, as broadly increased under antibiotic exposure (a total of 65 unique proteins). Conversely, proteins more abundant in N or AN, or less abundant in NA, were grouped as generally increased in non-antibiotic conditions (a total of 60 unique proteins). The relevant filtered pathway results are shown in **Fig. 4**. The unfiltered results are shown in **Figs. S13-S14**.

**Figure 4.**
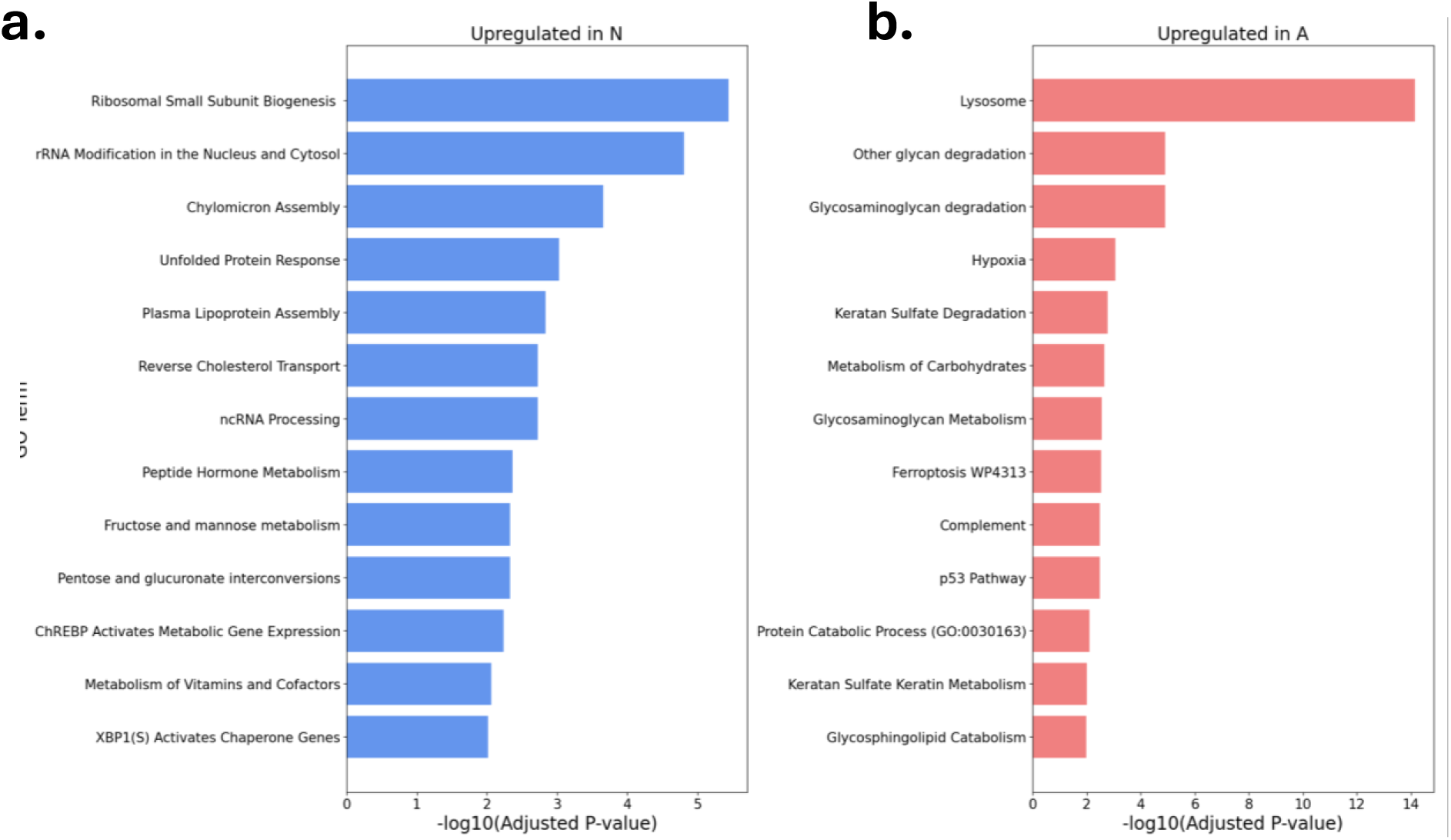
GO figures for terms upregulated in samples **a)** without or **b)** with antibiotics. Significant proteins were those with adjusted p-values ≤ 0.01. Redundant or irrelevant terms were manually removed. The unedited figures can be found in the Figs. S13,14. The genes set used were: MSigDB_Hallmark_2020, GO_Biological_Process_2023, Reactome_Pathways_2024, WikiPathways_2024_Human, and KEGG_2021_Human (accessed June 5, 2025).

Our original hypotheses were that antibiotics cause off-target effects on mitochondrial and ribosomal proteins. These plots reflect such results. Antibiotics may induce mitochondrial stress, resulting in alterations in metabolic pathways. This explains why terms like hypoxia are enriched in A, and various metabolic processes (*e.g.,* lipids, carbohydrates; Carbohydrate Response Element Binding Protein (ChREBP1 pathway)) are altered across both conditions (**Fig. 4**). This is consistent with known effects of aminoglycosides like streptomycin, which, in addition to targeting bacterial ribosomes, can bind eukaryotic mitochondrial and cytosolic ribosomes.^12^ We indeed found enrichment of ribosome biogenesis and rRNA terms in non-antibiotic-containing media (**Fig. 4a**). Thus, mitochondria (*via* altered metabolism) and ribosomal effects are reflected on the protein level.

Several additional pathway enrichments provided insight into broader cellular consequences of antibiotic exposure. For example, the increase in unfolded protein response, p53, complement, and ferroptosis pathways, which are enriched in antibiotic-conditions (**Fig. 4b**), may explain the observations regarding antibiotics’ effects on cell proliferation, cytotoxicity, and cancer.^32^

### Literature comparison

To contextualize our findings, we reviewed several relevant studies detailing the effects of antibiotics on mammalian cell cultures (**Table S2**). From this, **Table 1** highlights proteins and protein families that were differentially abundant in our study and at least one other relevant publication.

**Table 1.** Proteins of interest found to be differentially abundant in this study and at least one other relevant study. Underlined proteins/genes denote an exact match. Bolded families are prefixes for multiple proteins listed. An asterisk denotes unspecified protein match.

| Biological Relevance | Reference |  |  |  |  |  |  |
| --- | --- | --- | --- | --- | --- | --- | --- |
|  | This work | Ryu et al. | Mathieson et al. | Elliot et al. | Relier et al. | Kalghatgi et al. | Han et al. |
| Calcium homeostasis | <u>RGN</u> | <u>RGN</u> |  |  |  |  |  |
| Cell cycle/signaling | CDK16 |  |  |  |  |  | CDK2 |
|  | PPP1R2 | PPP2CB |  |  |  |  |  |
|  | TOP2B | TOP1 |  |  |  |  |  |
|  | <u>IGFBP1</u> | <u>IGFBP1,3</u> |  |  |  |  |  |
| Cytoskeleton | ACTG2, ACTR1A, PHACTR2, ENAH |  | ACT* | ACTB |  |  | ACTB |
|  | <u>KRT19,23,80</u> | <u>KRT23</u> |  |  |  |  |  |
|  | <u>AKAP12</u> | <u>AKAP12</u> |  |  |  |  |  |
| Glycolytic and ATP metabolic processes | <u>NME3,4</u> | <u>NME1</u> |  |  |  |  |  |
|  | PFKP |  |  | PFKM |  |  |  |
|  | <u>PKLR</u> |  | <u>PKLR</u> |  |  |  |  |
|  | AK1 | AK2 |  |  |  |  |  |
|  | ATP6V0A1 | ATP2B2 |  |  |  |  |  |
| Immune response | <u>ANXA3,4</u> |  |  |  | ANXA5 |  |  |
| Lipid metabolism | <u>PLIN2</u> | <u>PLIN2</u> |  |  |  |  |  |
|  | PLA2G15 | PLA2G12B |  |  |  |  |  |
|  | ACSL3 | **ACSS3,S5,L1,L5,M3** |  |  |  |  |  |
|  | <u>PECR</u> | <u>PECR</u> |  |  |  |  |  |
|  | <u>ABAT</u> | <u>ABAT</u> |  |  |  |  |  |
| Oxidative stress response | <u>ALDH1A1, L2, 4A1,18A1</u> |  |  |  | ALDH* |  |  |
|  | <u>GPX2</u> |  |  |  |  | GPX1 |  |
| Protein folding, chaperones, and proteostasis | <u>LRRC20</u> | <u>LRRC59</u> |  |  |  |  |  |
|  | <u>HSPB1, 90B1</u> | <u>HSPE1,A5,9,H1</u> | <u>HSP10,70,90</u> |  |  |  |  |
|  | <u>BAG2</u> | <u>BAG2</u> |  |  |  |  |  |
|  | <u>HYOU1</u> |  | <u>HYOU1</u> | HIF1a |  |  |  |
|  | UBAC1, UBFD1 | UBASH3B, UBE2Q1 | UBA1 |  |  |  | UBB |
|  | <u>NEDD4L</u> | <u>NEDD9</u> | <u>NEDD8</u> |  |  |  |  |
|  | <u>VCP</u> |  | <u>VCP</u> |  |  |  |  |
|  | <u>RNF149</u> | <u>RNF10</u> |  |  |  |  |  |
| Ribosome biogenesis | <u>UTP4,6,15,14A</u> | <u>UTP15,14A</u> |  |  |  |  |  |
|  | <u>RPF2</u> | <u>RPF2</u> |  |  |  |  |  |
|  | <u>WDR3,36,43</u> | <u>WDR77</u> |  |  |  |  |  |
| RNA processing | <u>PCBP2</u> |  |  |  |  |  | <u>PCBP2</u> |
|  | <u>DDX3,17,27,47, 52,54,56</u> |  |  |  |  |  | DDX5 |
|  | <u>RBM22,28</u> | <u>RBM24</u> |  |  |  |  |  |
| Translation | <u>EIF6,2AK4,4H</u> | <u>EIF2B2,3J</u> |  |  |  |  |  |
|  | <u>SRPRB</u> | <u>SRPRB</u> |  |  |  |  |  |
| Transmembrane transport | <u>SLC3A2,39A4, 10,14,O4C1</u> | <u>SLC6A14,20A1,7A11, 1A4,35B1,22A7,6A4, 39A4,38A4</u> |  | <u>SLC2A1,3</u> |  |  |  |
| Xenobiotic response | <u>ABCB1</u> | <u>ABCB1,C6P1</u> |  |  |  |  |  |
| Transcription regulation | ZFAND3, ZNF281 | ZFP36L |  |  |  |  |  |

In particular, our study follows a similar design to that of Ryu *et al*.^3^ Ryu *et al.* used HepG2 cells grown from passage 37 under the same antibiotic- and non-antibiotic media used here, except in a single 6-well plate (3 replicates) for two passages (21 days). Using RNA-seq and ChIP-seq, the authors found 209 genes that were differentially expressed between A and N. While our study differs in sample size and exposure time, the results of their transcriptomics studies allow for comparison with our proteomic results (**Fig. 5**, **Table 1**). Of the 9,964 genes in the Ryu *et al*. dataset annotated with UniProt IDs, we extracted 206 differentially expressed genes.

**Figure 5.**
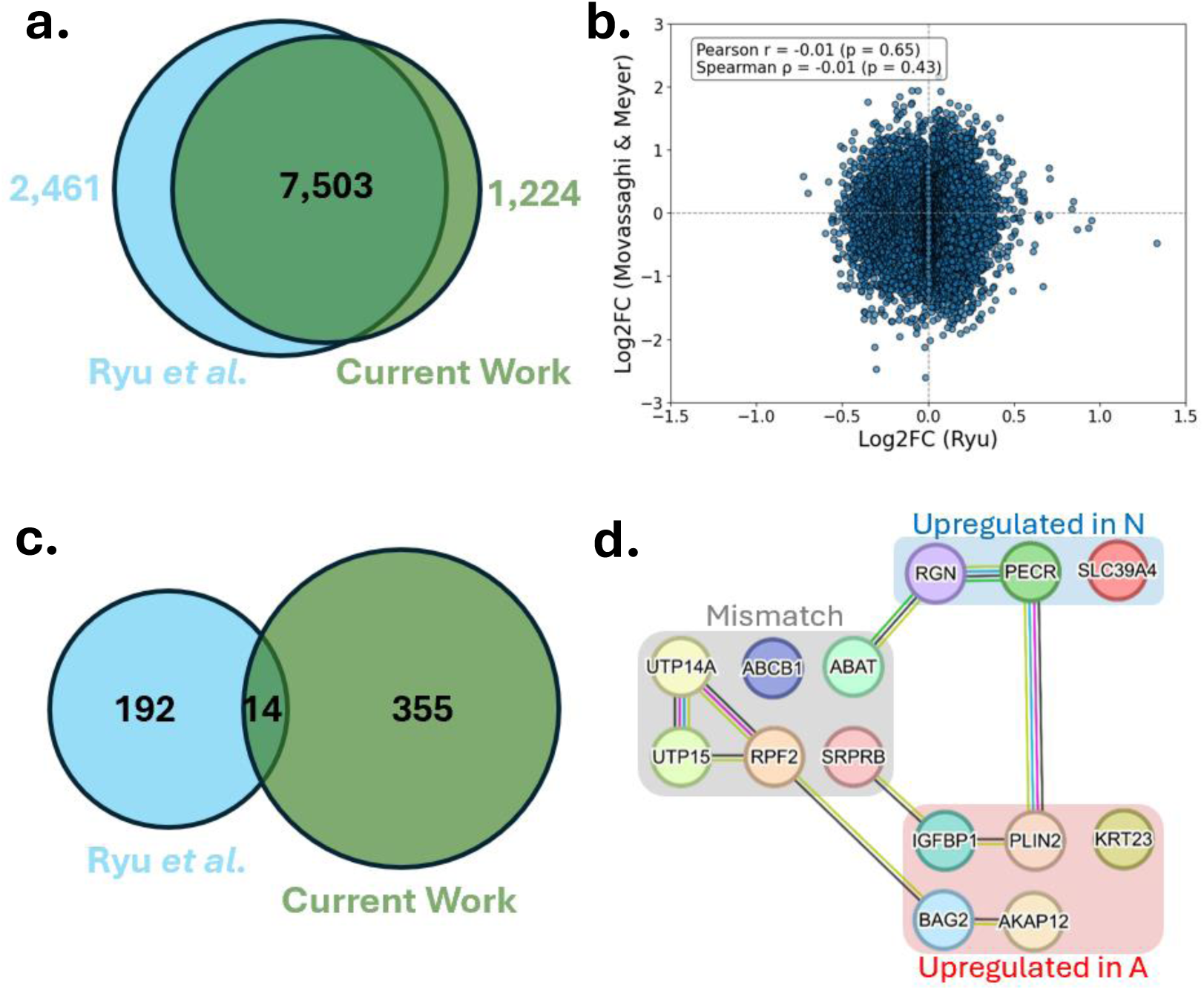
Literature comparison. **a)** Overlap of proteins detected in this work versus gene names (with UniProt IDs) reported by Ryu *et al*. **b)** Fold-change (log2 scale) correlation between this work and Ryu *et al.* Both fold changes are from the second passage only. **c)** Overlap of significant proteins. A threshold of p ≤ 0.1 was used to match the report by Ryu *et al*. **d)** STRING diagram of the overlapping proteins from c.

Overall, we found substantial overlap in coverage of our proteomic dataset with the transcriptomic dataset of Ryu *et al.*; after filtering for redundant isoforms, the overlap was nearly 80% (7,503) (**Fig. 5a**). We then compared the log2 fold changes reported by Ryu *et al.*(from their second passage) with our log2 fold changes from our second passage, focusing only on treatment groups N *versus* A. The results were not correlated between studies (**Fig. 5b**). To test whether this discrepancy was due to differences in exposure duration rather than passage number, we performed an additional comparison using our sixth passage (19 days), which more closely matched the 21-day exposure in Ryu *et al*. This analysis yielded a small, but still statistically insignificant, positive correlation (r = 0.02, **Fig. S15**). This lack of correlation may be expected, as transcriptomics and proteomics are known to be poorly correlated,^33,34^ especially under oxidative stress.^35^ Even within the report of Ryu *et al.,* the correlation between ChIP-seq and RNA-seq was as low as 0.15, underscoring the complexity of gene regulatory responses to antibiotics at different biological levels.

We then compared our differentially abundant proteins from the LME model with those defined as differentially expressed by Ryu *et al.* (**Fig. 5c**). To match their adjusted *p*-value threshold, we reduced our adjusted *p*-value cut-off to <0.1 and identified a total of 383 differentially abundant proteins (208 upregulated and 175 downregulated). Ryu *et al*. identified a total of 209 differentially expressed genes (157 upregulated and 52 downregulated). Of these, 14 exact matches (by UniProt ID) were found (**Fig. 5d**). Five proteins were significantly upregulated in A in both data sets, and three proteins were significantly upregulated in N in both datasets. However, six proteins were downregulated in Ryu *et al.* but upregulated in our data. No proteins downregulated in our data were reported as upregulated in Ryu *et al*.

A reason for this mismatch could be due to differences in passage numbers, passage-associated drift, experimental design, or the well known discordance between proteomes and transcriptomes. Our work utilized HepG2 cells at passage 22 with a longitudinal crossover design, whereas Ryu et al. employed cells at passage 37, which were collected after two passages, for two treatment conditions. Noted genetic differences occur in immortalized cell lines across passages, especially at high passage numbers.^18^ Furthermore, it was unclear whether the previous passages in Ryu *et al.* utilized antibiotic- or non-antibiotic-containing media. This shift in treatment baseline can result in contradictory findings, as suggested by Elliot *et al*.^8^

To broaden the search for similarities, we examined matches within the same isoform or protein family (**Table 1**), as well as similarly enriched pathways, across a survey of relevant studies. For example, we found CPT1A to be differentially abundant, while Ryu *et al.* found dysregulated mitochondrial carnitine shuttle pathways. CPT1A is the rate-limiting enzyme within the carnitine shuttle pathway, essential for fatty acid oxidation, which β-lactams are known to inhibit.^36^ Similarly, Ryu *et al.* reported dysregulation of forkhead box (FOX) transcription factors, which can bind with XBP1, an unfolded protein response (UPR) pathway enriched in our dataset.^37^ Other UPR pathways include eukaryotic translation initiation factors (EIFs), found in both our dataset and Ryu *et al*. The ubiquitin/proteasome protein degradation pathway is also activated as proteins accumulate, and protein translocation in the endoplasmic reticulum (ER) is mediated by ribosomal-binding SEC proteins (SEC11, 31, and 61 were all increased in N in our data).^38^ This can increase ER stress, leading to lipid accumulation in hepatocytes. Excess lipids can be transiently stored as lipid droplets. Lipid droplets can then undergo lysosome-dependent autophagy, which is reflected by our findings of differential abundance in alpha beta hydrolase domain (ABHD), RAB, heat-shock (HSP) and perilipin (PLIN) proteins. Accordingly, we also identified differentially abundant lysosomal proteins involved in lipid metabolism, including hexosaminidase B (HEXB), phospholipase A2 (PLA2G15), and cathepsin C (CTSC). These pathways explain the robust changes in proteins related to lipid metabolism, ER stress, chaperones, and protein turnover across studies (**Table 1**, **Fig. 6**). They may be a downstream effect of altered ribosomal or mitochondrial function.

**Figure 6.**
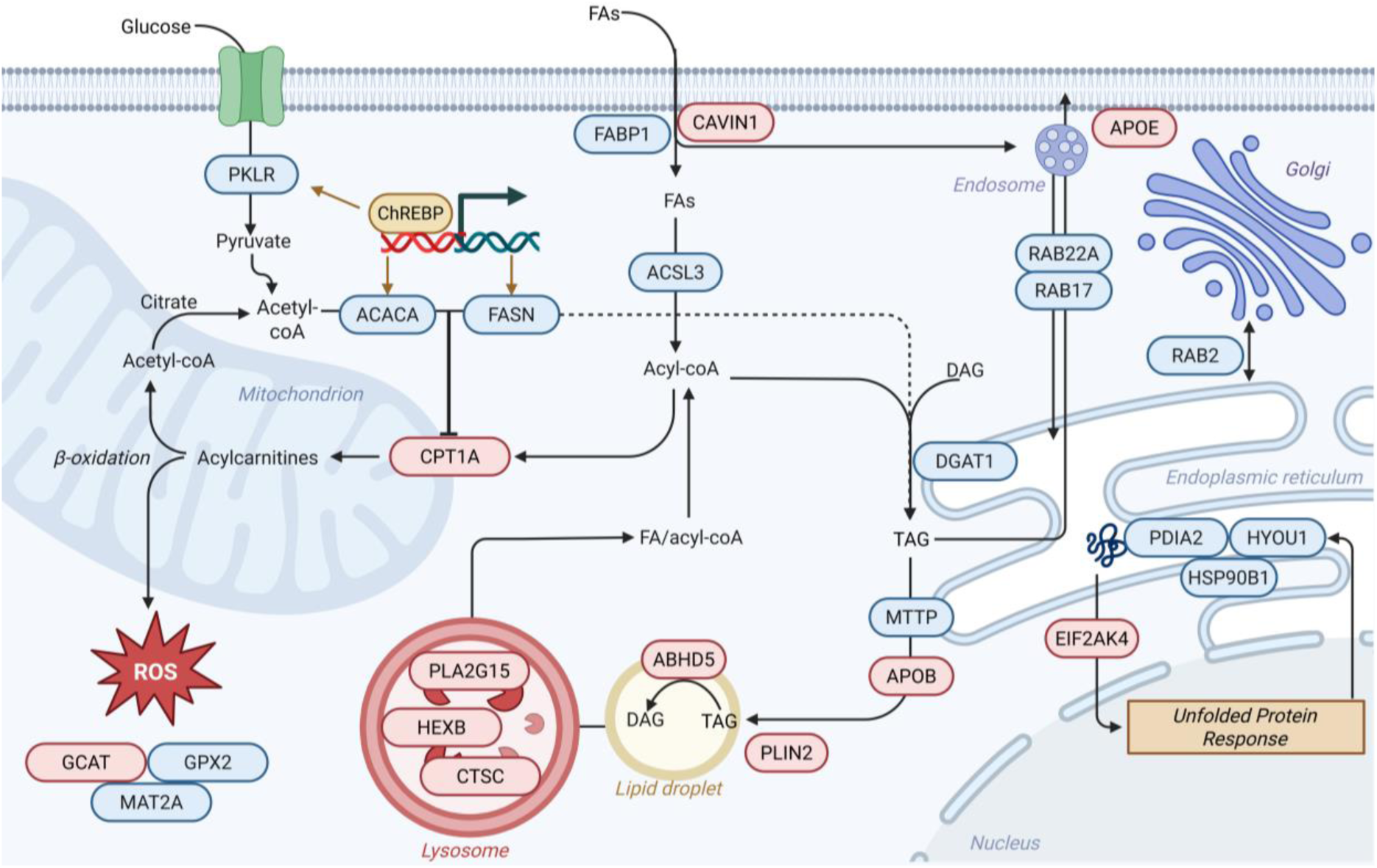
Possible dysregulated protein pathways involved in lipid transport and metabolism for HepG2 cells treated with or without antibiotics. Red UniProt IDs represent proteins upregulated in antibiotics, and blue represents proteins upregulated in non-antibiotics. All proteins shown were differentially abundant in this work.

Other, smaller-scale studies using various antibiotics and cell lines also provide insight into conserved biological changes. Llobet *et al*. demonstrated that the use of penicillin, streptomycin, amphotericin (AmB), and gentamicin influences the differentiation of human adipose tissue-derived stem cells into adipocytes.^9^ They found a 40% increase in reactive oxygen species (ROS) production in the presence of antibiotics, as measured by the rise in hydrogen peroxide production. When treated with the antioxidant N-acetyl-L-cysteine (NAC) in addition to antibiotics, intracellular lipid droplets and leptin staining decreased by up to 70%, compared to increases of up to 36% and 290%, respectively, in PenStrep. Interestingly, in contrast with our and Ryu *et al.* results, authors did not find significant effects on mitochondrial enzymes using colorimetric assays.

Kalghatgi *et al.* explored aminoglycosides and β-lactams in relation to mitochondrial dysfunction and oxidative damage *in vitro* (using MCF10A cells) and *in vivo* (in mice).^39^ The authors found that clinically relevant treatments caused mitochondrial dysfunction and the overproduction of ROS. For example, glutathione peroxidase (GPX1), among other antioxidant defense genes, was upregulated up to 10-fold after prolonged antibiotic treatment. Interestingly, these effects were also alleviated by the addition of NAC. Of these, in our work, we found that GPX2 was higher in cells without antibiotics.

Relier *et al.* examined the effect of PenStrep on the phenotypes of cancer cells.^40^ They found these antibiotics inhibited sphere-forming ability in six cancer cell lines across three different tissue types. Staining for annexin V (ANXA5) revealed no significant differences after PenStrep treatment, while our results showed increased ANXA isoforms in antibiotic-containing samples. Meanwhile, staining for aldehyde dehydrogenase (ALDH) showed a substantial decrease in PenStrep, which is in agreement with the ALDH isoforms detected in our dataset.

Elliot *et al.* investigated the effects of gentamicin in three mammary cell lines.^8^ This study showed a significant upregulation of hypoxia-inducible factor 1 (HIF1a), phosphofructokinase (PFK-M), lactate dehydrogenase (LDH-A), and glucose transporters SLC2A1 and SLC2A3 in gentamicin-treated relative to controls. Gentamicin was also found to significantly increase mitochondrial superoxide production within 24 hrs. These results are in general agreement that solute carrier family (SLC) proteins, as well as glycolysis and ROS pathways, have conserved effects for antibiotics beyond PenStrep.

Mathieson *et al.* used MCF-7 breast cancer cells with PenStrep, AmB, and control.^6^ AmB is a commonly used antifungal agent and is often combined with PenStrep to form an antimicrobial cocktail (*i.e.,* “anti-anti”). They demonstrated that no differences in contamination, dead cells, or cell cycle phase that would account for between-culture variations in confluence. Instead, they reported widespread decreases in chaperone, cytoskeletal, and ER stress-related proteins in PenStrep-treated cells. In our dataset, hypoxia upregulated 1 protein (HYOU1), HSP90, valosin containing protein (VCP), and pyruvate kinase (PKLR) were similarly decreased, further supporting a conserved stress-response proteome.

Han *et al.* reported the dose-dependent effects of gentamicin, streptomycin, and penicillin on mouse blastocysts using RNA-seq.^41^ For the lowest doses tested, roughly 1,800 genes were significantly differentially expressed. For penicillin and streptomycin specifically, the top KEGG pathways affected included p53, RNA splicing and transport, ubiquitin-mediated proteolysis, cholesterol and lipid metabolism, and glycan biosynthesis, which align with pathway results from our work (**Fig. 4**).

Based on the above literature, summarized in **Table 1**, a consistent alteration in lipid metabolism was observed. Other conserved effects across studies include protein turnover, cytoskeletal maintenance, oxidative stress, ribosome biogenesis, and membrane transporters. A list of possible protein pathways dysregulated in lipid metabolism is shown in **Figure 6**. All proteins/pathways shown were differentially abundant in our data.

For example, the carbohydrate response element binding protein (ChREBP) pathway was enriched in N (**Fig. 4**). ChREBP is a transcription factor that regulates glycolytic and lipogenic proteins such as pyruvate kinase (PKLR), acetyl-coA carboxylase alpha (ACACA) and fatty acid synthase (FASN),^37^ all of which are more abundant in N. The latter proteins are involved in *de novo* lipogenesis, where ACACA produces malonyl-CoA, followed by the rate-limiting production of palmitate by FASN. Malonyl-CoA is a known inhibitor of CPT1A, preventing long-chain fatty acid oxidation.^37^ Meanwhile, in A, CPT1A and thus fatty acid oxidation are upregulated, generating ROS. The liver is a key regulator of glutathione, which protects against ROS, and also has roles in detoxication of xenobiotics.^20^ This may account for the dysregulation of various proteins involved in glutathione synthesis *via* amino acid metabolism (GCAT, MAT2A) or antioxidant response (GPX2), the latter being ferroptosis-related (a process upregulated in A). However, glutathione synthesis is rate-limited by the availability of cysteine and the activity of glutamate-cysteine ligase. The latter accounts for MAT2A, which is involved in the methionine cycle to produce cysteine. Some evidence suggests that glutathione synthesis may be jointly regulated by both glycine and cysteine.^42^

The liver is highly metabolic, and principally utilizes fatty acids. Free fatty acids enter liver cells *via* diffusion or active transport by proteins like CD36 and are eliminated via oxidation or exocytosis of lipoproteins. Meanwhile, cholesterol and lipoproteins enter through receptor-mediated endocytosis. These processes are heavily regulated to maintain a steady state and prevent accumulation. In HepG2 cells, fatty acid binding protein 1 (FABP1) and CAVIN1 are involved in the transport of fatty acids.^37,38^ FABP1 is also found in the cytosol, where it binds various lipid substrates. Acyl-CoA synthetase long-chain (ACSL) enzymes then produce fatty acyl-CoA. ACSL1 is found on the outer mitochondrial membrane and interacts with CPT1A in hepatocytes. Both ACSL1 and ACSL5 were identified by Ryu *et al.* and are known to be highly expressed in liver^37^, while we identified the upregulated ACSL3 isoform in the absence of antibiotics. ACSL3 is expressed on lipid droplets and is involved in de novo lipogenesis. Meanwhile, acyl-CoA thioesterase 2 (ACOT2) catalyzes the reverse reaction of ACSL isoforms, leading to lipid catabolism, and was upregulated in N. The coordinated regulation of ACOT and ACSL remains under study.^38^

During triglyceride synthesis, diacylglycerol acyltransferase (DGAT) transforms diacylglycerols (DAGs) to triacylglycerols (TAGs) in the ER, which can then form lipid droplets.^38^ These droplets can then undergo autophagy with a lysosome or be secreted as very-low-density lipoprotein (VLDL). In lipid droplets, the perilipin (PLIN) superfamily and microsomal triglyceride transfer protein (MTTP) incorporate TAGs into apolipoprotein B (APOB), which move from the ER to Golgi to form VLDL.^38^ The transport of VLDL also accounts for the differential abundance of several vesicle trafficking proteins (RAB2B,17,22A and SEC11C,31A,61B) found to be increased in N.

Overall, the lipid metabolism signaling in our data appears mixed, with some proteins indicating more catabolism and others indicating more storage. A possible source of error is differences in cell density.^43^ However, several precautions were taken to avoid such effects, including matched cell seeding densities, visual confirmation of matched confluency throughout the experiment, and the use of linear mixed-effect models to identify consistent effects across passages. Future work may focus on elucidating the precise mechanisms underlying these differences.

## CONCLUSIONS

Despite its ubiquity, we demonstrate that supplementing mammalian cell culture media with antibiotics yields a range of ribosomal and mitochondrial variations in the proteome. The breadth of our results enabled comparisons across protein, transcript, and pathway-level analyses. While we observe some conserved agreements related mainly to lipid metabolism, oxidative stress, and protein turnover, the magnitude and direction of many effects appear to be dependent on cell line, culture conditions, concentrations, time (including passage), as well as the type of antibiotic used. In agreement with previous transcriptomics advice, proteomic studies (especially those involving *in vitro* models of metabolism) should consider the confounding effects of antibiotic supplementation. In particular, for the bulk proteomic analysis of HepG2 cells, care should be taken when 1) comparing results across studies using antibiotics versus not, or using different types of antibiotics; and 2) if treating acutely with antibiotics, cell cultures should be re-conditioned in control media for at least three passages to ensure a more consistent proteomic profile. Lastly, a potential area for future work is the use of antioxidants, such as N-acetyl-L-cysteine, to mitigate off-target effects involving reactive oxygen species, in conjunction with antibiotics.

## METHODS

### Materials

The following materials were obtained from ThermoFisher Scientific (Waltham, MA): Dulbecco’s Modified Eagle Medium (DMEM; 11995-065); phosphate buffered saline (PBS; 14190144); dialyzed fetal bovine serum (FBS; A3382001); penicillin-streptomycin (PenStrep; 15-140-122); radioimmunoprecipitation assay buffer (RIPA; AAJ62524AE); ammonium bicarbonate (AC393210010); 0.25% Trypsin-EDTA (25-200-056); 1M Tris-HCl pH 8 (15568025); LC-MS grade 99% formic acid (A117-50); LC-MS grade 0.1% formic acid in water (v/v) (LS118-1). The following materials were obtained from Millipore Sigma (Burlington, MA): Roche cOmplete protease inhibitor cocktail mini tablets (11836153001); HPLC grade ethanol (59828-1L); anhydrous calcium chloride (CaCl_2_, C5670-100G); HPLC-grade acetonitrile (34851); Pierce HeLa protein digest standards (88329); dithiothreitol (DTT) and 3-indoleacetic acid (IAA). Sequencing-grade modified trypsin (V5113) was purchased from Promega (Madison, WI). Silica magnetic beads (786-916) were purchased from G-Biosciences (St. Louis, MO).

### Cell culture

HepG2 cells were a gift from Shelly Lu’s lab. The HepG2 cells were cultured in high-glucose DMEM with 10% fetal bovine serum (FBS) and antibiotics (1% PenStrep; 100 U/mL penicillin, 100 µg/mL streptomycin) for five passages to ensure the absence of latent microbial contamination. All incubation occurred at 37 °C and 5% CO_2_. At passage 22, one confluent T-75 flask of HepG2 cells was split into two six-well tissue-culture-treated plates. Each well of each plate contained a seeding density of roughly 0.2×10^6^ cells. One plate contained cells in high-glucose DMEM with 10% FBS and 1% PenStrep (antibiotic-containing media; A), and the other plate contained cells in high-glucose DMEM with 10% FBS (no antibiotics; N). Media changes were performed approximately every two days. After approximately 72 hours, roughly equal confluency of > 80% (∼1.2×10^6^ cells) was confirmed across plates using an inverted microscope. One well of each plate was used to seed the six-well plate for the following passage; *e.g.,* one well of A_1_ was used to seed the entire plate of A_2_. This was done by placing 1 mL of 0.25% trypsin-EDTA in the well, incubating for 3 min, diluting with serum-free DMEM, centrifugation at 1,000 rpm for 5 min, aspirating, and resuspending the pellet in 12 mL of the corresponding treatment media. The remaining five wells were rinsed with PBS and lysed using 125 µL of RIPA with a dissolved protease inhibitor cocktail tablet (1 tablet/10 mL), and then placed on ice for 5 min. Each well was then scraped and transferred to Eppendorf tubes. The tubes were centrifuged at 14,000 rpm for 15 min, sonicated for 30 s, and the supernatant was transferred to a new Eppendorf tube. Samples were stored in -80 °C before sample preparation. This process was repeated for passages 1-5. On passage 6, to split into cross-over conditions, two wells were harvested per plate, while the remaining four were lysed as before. One well was resuspended in antibiotic-containing media, and the other in non-antibiotic-containing media. This resulted in four total plates from passage six onwards: A_6_, A_5_N_1_, N_6_, N_5_A_1_. The harvest and lyse protocol was then performed as described for passages 1-5 for the remaining passages up until passage 9. All cultures were checked roughly every 24 hr for any evidence of microbial contamination using an inverted microscope.

### Sample preparation

All samples were prepared on the same day using an automated single-pot, solid-phase enhanced sample preparation protocol (SP3)^17^ using a KingFisher Flex (ThermoFisher). Samples were thawed and 5.8 µL of lysate was transferred into each well (∼ 33 µg protein/well). Including quality control samples, approximately 160 samples were prepared across two 96-well plates. The quality controls included a pooled sample (5 µL of all samples mixed together) and a blank (containing only RIPA buffer; no cell lysate). Samples were reduced by incubation with DTT for 30 min at RT, followed by alkylation with IAA for 30 min at RT in the dark. Next, 6.6 µL of paramagnetic silica beads were added to achieve a 1:10 ratio of protein: beads (w/w). Beads were washed with MilliQ water using the KingFisher flex prior to sample addition. Next, 105 µL of acetonitrile (LC-MS grade) was added to each well to achieve a final concentration of 70% acetonitrile (v/v) in each well. The samples were incubated at RT for 18 min with gentle agitation. Next, the samples with beads containing precipitated protein were washed twice in 500 µL 80% ethanol, followed by two washes in 500 µL 100% acetonitrile. The beads were then transferred into a 50 mM solution of Tris-HCl containing 10 mM CaCl_2_ and 1:25 protein: trypsin(w/w). Samples were covered and incubated with gentle agitation at 37 °C overnight. The next day, the beads were removed from the samples and the samples were diluted with 5.5 µL of 10% formic acid followed by 109 µL of 0.1% formic acid. The plates were then centrifuged and transferred to a new 96 well plate for analysis.

#### LC-MS

Samples were organized into a blocking pattern containing one sample from each treatment and passage. The injection order was randomized within each block. A blank and pooled sample was run before and after each block. The performance of the system was tested using injections of solvent blanks and HeLa protein digest standards. An injection volume of 10 µL was used. A Vanquish Neo LC was used with PepSep C18 15 cm x 150 µm, 1.5 µm at 55 °C (Bruker Corporation, Billerica, MA). A 30-minute chromatography gradient was used, using 0.1% formic acid in water (solvent A) and 80% acetonitrile with 0.1% formic acid in water (solvent B). The starting composition of solvent B was 4%, increasing to 9% at 2 min, 25% at 18 min, 35% at 27 min before column wash and equilibration. A pump curve value of 5 and flow rate of 1.2 µL/min was used for all gradient steps. A Thermo Orbitrap Astral was used for mass spectrometry. Data were acquired using data-independent acquisition with the Orbitrap resolution set to 240,000 and a 3 *m/z* isolation window spanning 380-980 *m/z*.

### Data analysis

DIANN 1.9 was used to quantitate proteins from the raw data in library-free mode. A human reference proteome .fasta file (downloaded January 27, 2025) containing canonical proteins and isoforms (83,062 proteins, and 20,562 genes) was used to generate an *in silico* library. Three missed cleavages were allowed. All other settings were default. The output pg_matrix from DIANN was analyzed in Jupyter notebooks using Python version 3.12.2, numpy version 1.26.4, pandas version 2.2.2, and anndata version 0.10.9. Plots were made using matplotlib 3.8.4 and seaborn 0.13.2. Pre-processing, imputation, and principal components analyses (PCA) were done using scanpy version 1.10.2 and scikit-learn version 1.4.2. Prior to PCA analysis, the data was scaled across both the samples and proteins. After removing blanks, quality control samples, and 7 sample outliers from the dataset, any proteins present in less than 50% of samples were removed. This resulted in 9,518 proteins out of 10,255 (93%) proteins. All remaining 119 samples contained at least 8,924 proteins (<7% missingness), up to 9,374 proteins in the highest sample (average±standard deviation of 9,210±106 proteins). Samples were log-transformed then normalized by the total protein quantity present prior to imputation of missing values using a *k*-nearest neighbors imputer with five nearest neighbors. Each sample was then annotated by plate (*i.e.,* 1-2), replicate (*i.e.,* 1-5), passage (*i.e.,* 1-9), and condition (*i.e.,* A, N, AN, or NA). Statistical tests such as t-tests were done using scipy stats package version 1.13.1. Multiple tests correction was performed using Benjamini-Hochberg correction and the adjusted p-value significance threshold was set to a p ≤ 0.01 cutoff, unless otherwise noted. Linear mixed effect models utilized statsmodels 0.14.2. The gseapy package version 1.1.4 was used for gene set enrichment analyses. All STRING diagrams were created using string-db.org (version 12.0) using the full network type, with evidence-based edges, all active interaction sources, minimum low confidence interaction scores (0.150), and no interactors.

## Acknowledgements

We thank for Shelly Lu lab for gifting us a stock vial of HepG2 cells and for comments on the manuscript. We thank Bora Onat and Jesus de Jose Munoz Estrada for assistance with cell culture, and Simion Kreimer for help in operating the Orbitrap Astral. Biorender.com is acknowledged for generation of the table of contents graphic and Figures 1, 6.

## Data and code availability

All raw data and DIANN output files can be accessed through MassIVE under dataset identifier MSV000098243 (doi:10.25345/C59P2WK1T). All analysis code can be accessed at https://github.com/xomicsdatascience/hepg2_abc.

## Contributions

CSM and JGM conceived of the study. CSM performed the experiments and analyzed the data. JGM supervised the project. All authors wrote and approved of the manuscript.

