## Supplementary figures for "Antibiotics Rewire Core Metabolic and Ribosomal Programs in Mammalian Cells"

Supplementary data files:

- Spreadsheet 1: full linear mixed effect model results

**Supplementary Note 1**

We examined the total protein quantities annotated by various experimental factors. Throughout, A refers to all samples in all passages exposed only to antibiotic-containing media ($A_{1-9}$), N refers to all samples in all passages exposed to only non-antibiotic containing media ($N_{1-9}$), while AN and NA refer to the crossover conditions present only in passages 6-9, i.e., $A_5N_{6-9}$ and $N_5A_{6-9}$. We observed negligible differences in total protein content by the sample preparation plate (*e.g.*, P1 *vs*. P2; two separate 96-well plates were used for sample preparation and analysis of the 119 samples), indicating a lack of batch effects (**Fig. S1a**). We also observed little difference in total protein content related to treatment condition (**Fig. S1b**), indicating consistent confluency and recovery regardless of antibiotic supplementation. However, when examining total protein quantities across passages, passages five and six were slightly, but consistently, lower compared to the remaining passages (**Fig. S1c**).

To identify the samples and sources causing this variation, we examined the scores plot of the first two principal components of the data, annotated by passage number (**Fig. S1d**). We noticed that one sample from passage four, 75% of the samples from passage five, and all of the samples from passage six were grouping separately along the PC1 axis from the remaining passages. This effect was not specific to a given plate or condition (**Fig. S2**). To identify what proteins were contributing to this distinguishing variation in passages 5 and 6, we examined the top loadings contributing to PC1 (**Fig. S1d**). A pre-ranked gene set enrichment analysis (GSEA) of these loadings revealed a possible cell-cycle dependent effect (terms G2-M checkpoint and E2F targets) along PC1 (**Fig. S1e**). These two terms were enriched in separate directions from the other major terms related to fatty acid metabolism, oxidative phosphorylation, and unfolded protein response. The latter terms are expected to occur given the experimental design and have been previously shown in transcriptomic data of HepG2 cells with and without antibiotic treatment.[1] However, cell cycle differences are less well characterized, but were statistically insignificant across antimicrobial treatments in MCF-7 breast cancer cells.[2] We thus hypothesized that proteins related to the cell cycle were responsible for the variation in the positive direction of PC1, suggesting that passages 5 and 6 may have been harvested at sub-confluent stages relative to the remaining passages.

Indeed, during the passaging and harvesting of passage six, these plates were noted as slightly less confluent when checked under the inverted microscope. Given that 75% of all conditions from passage five were skewed towards the more positive axis of PC1, along with 100% of passage six, we hypothesize this is simply a difference in culture confluency captured along PC1. Likely, passage five was harvested at a slightly sub-confluent time point, resulting in a marginally lower seeding density for passage six, which then corrects by passage seven. When we re-analyzed the dataset after excluding data from passages five and six, the loadings terms related to G2-M and E2F targets were no longer among the top loadings, while terms more likely to be associated with the experimental design appeared (*e.g.*, bile acid and xenobiotic metabolism, adipogenesis; **Fig. S3**). Other results related to differential expression and pathway analysis presented below were also minimally affected, regardless of whether or not we excluded these passages. For example, slightly fewer differentially expressed proteins were found when these passages were excluded, likely reflecting the loss in statistical power due to the smaller sample size; conclusions ascertained from the pathway analyses remained the same (**Figs. S4-S5**). For these reasons, we included data from all passages in the remaining analyses and accounted for passage-by-passage variation using a linear mixed-effects model (*vide infra*).

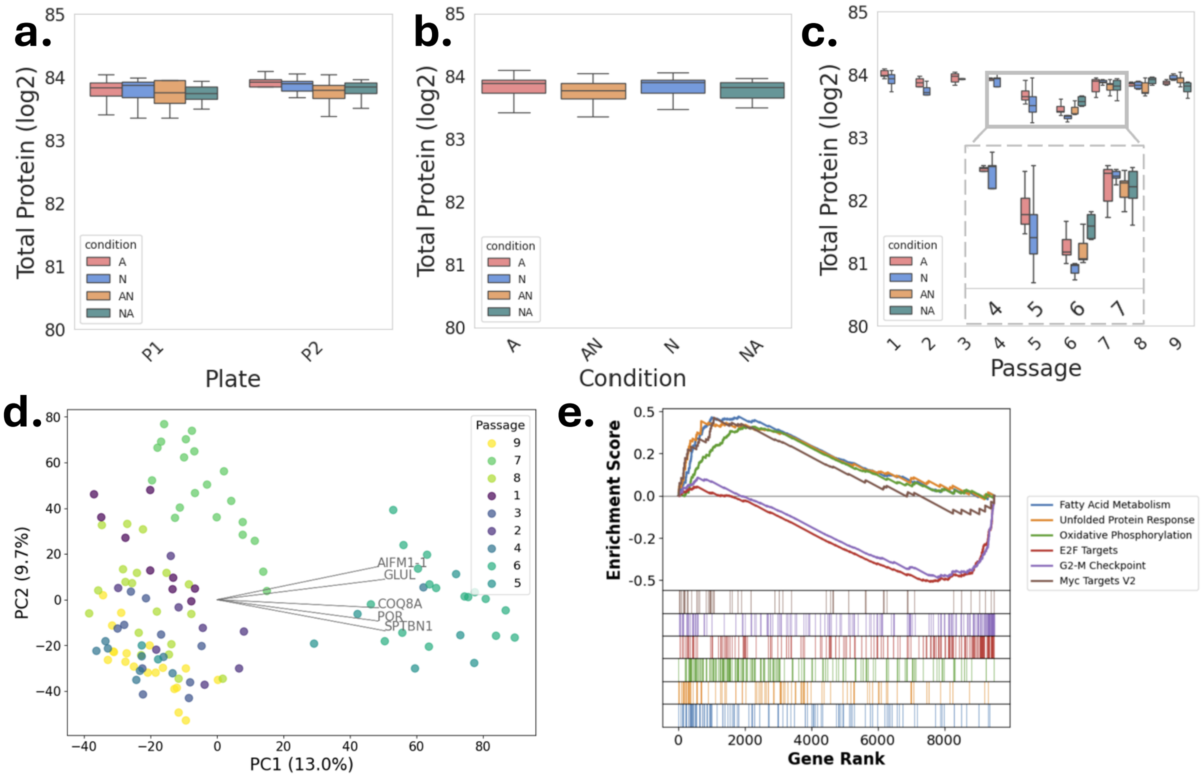


**Figure S1. Global differences across conditions.** Boxplots of total protein quantities by **a)** plate, **b)** condition, and **c)** passage (inset is zoomed in on the y axis). **d)** Principal components scores plot of samples annotated by passage. Gray lines represent the top five proteins of the PC1 loadings. **e)** The top six terms from the gene set enrichment analysis of the PC1 loadings using the MSigDB_Hallmark_2020 gene set.

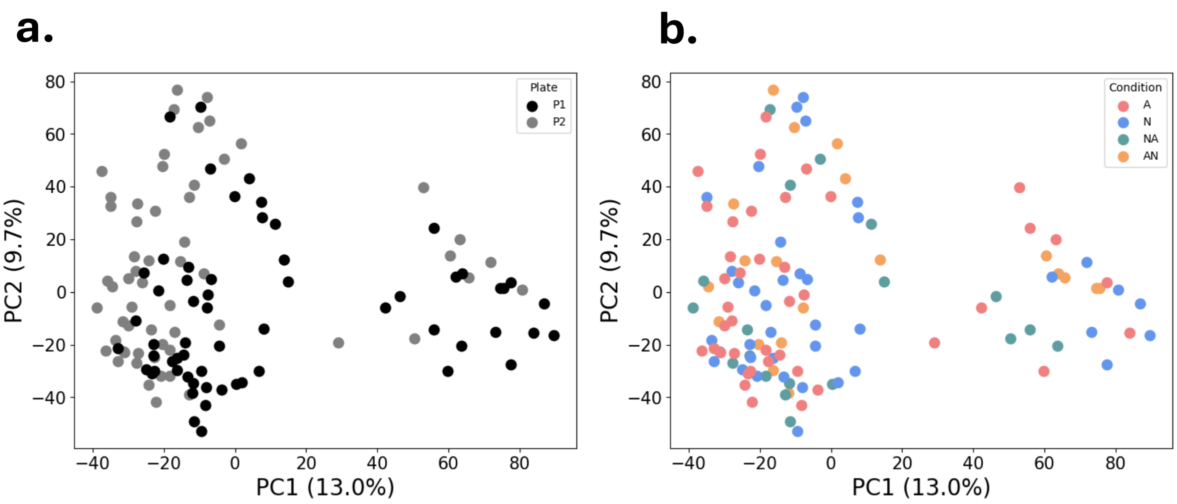


**Figure S2.** Principal components scores plot annotated by **a)** plate and **b)** condition.

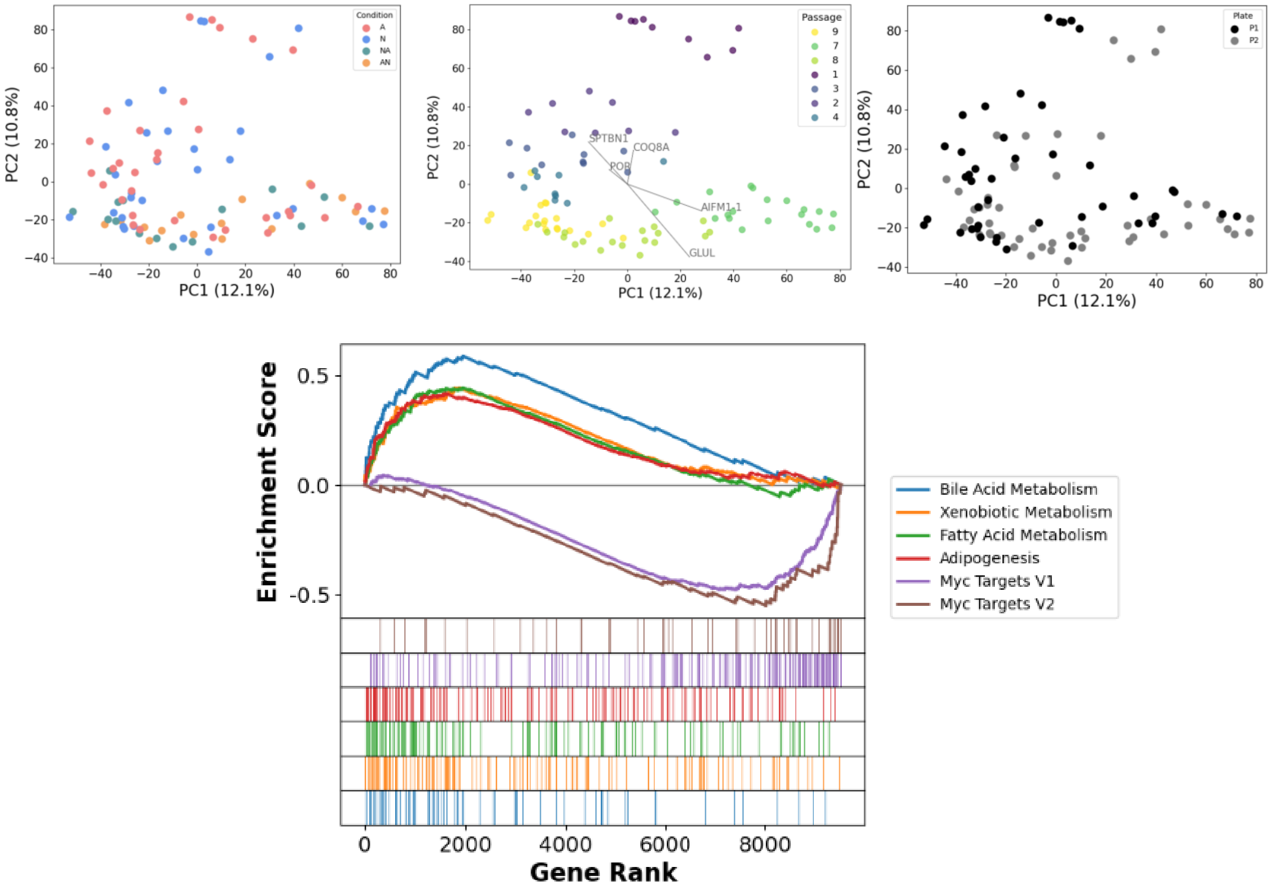


**Figure S3.** Recreated panels from Figs. S1, S2 with passages 4 and 5 excluded.

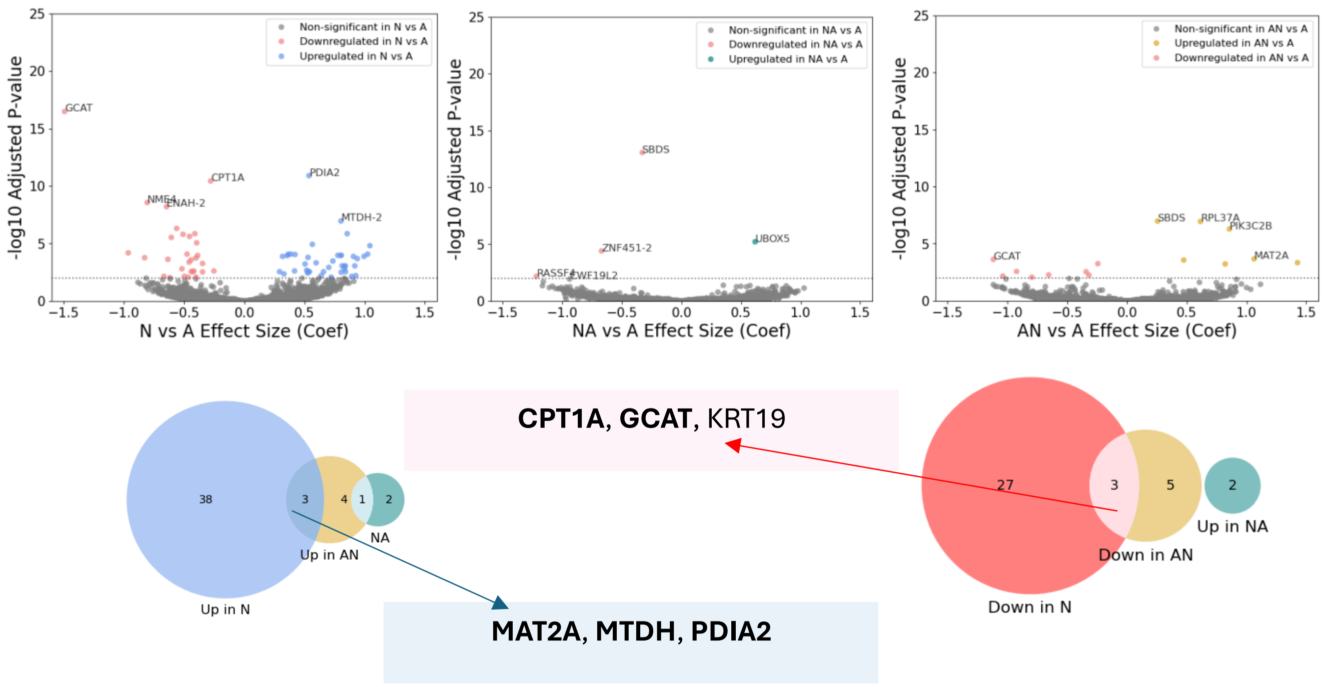


**Figure S4.** Recreated panels from Fig. 3 from the main text with passages 4 and 5 excluded. Bolded UniProt IDs represent exact matches with Fig. 3 proteins.

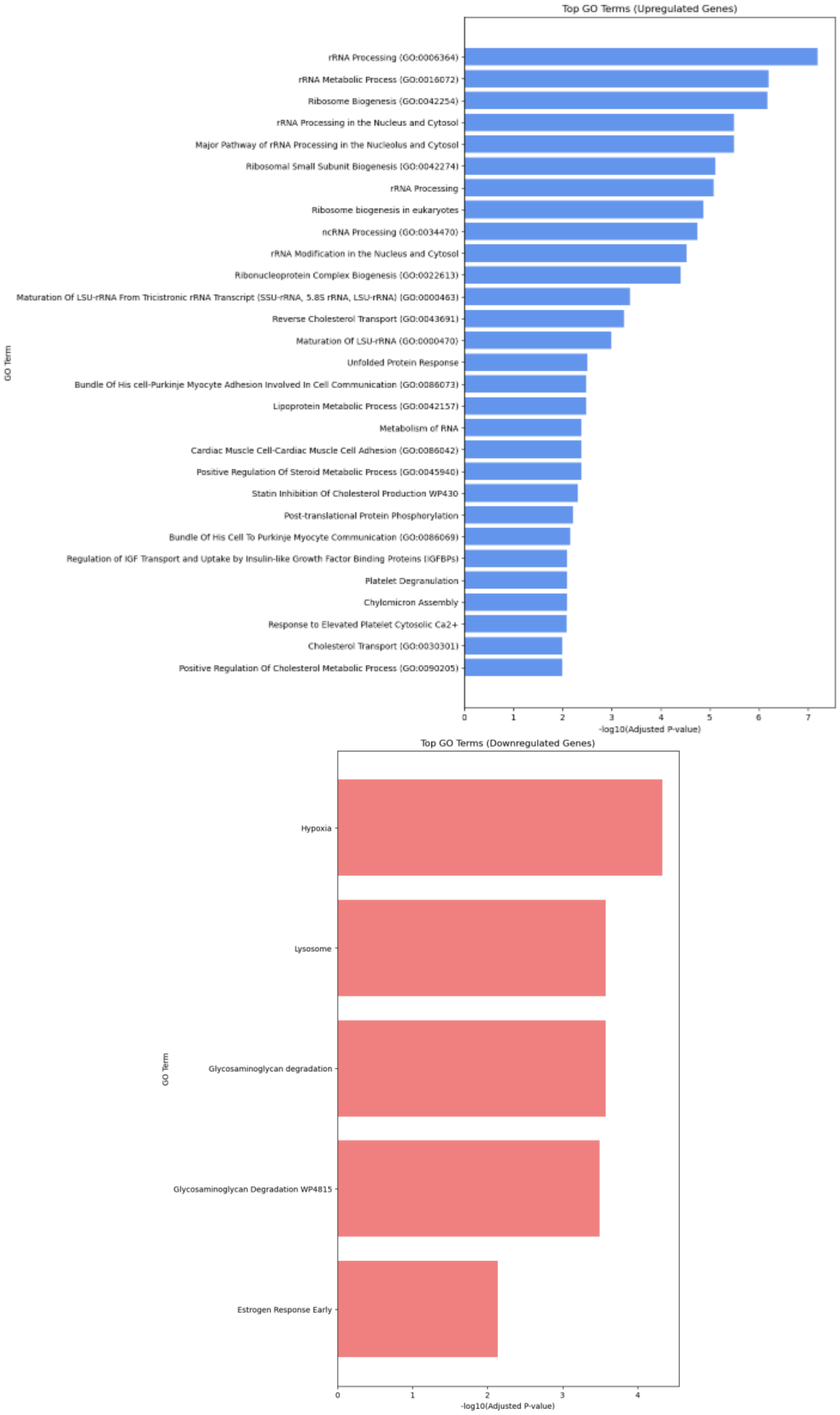


**Figure S5.** Recreated Fig. 4 from the main text with passages 4 and 5 excluded.

**Supplementary Note 2**

For comparison, we aggregated all A and N passages (1-9), performed a *t*-test, and found six differentially expressed proteins (**Fig. S6**). However, this analysis does not account for passage-by-passage variation, which appears to be a significant source of variation compared to condition (**Fig. S1a-d**). It also ignores the cross-over conditions (NA, AN). We then used multiple *t*-tests to compare all pairwise conditions, separated by passage (**Figs. S7, S8**). However, this approach does not account for correlation across passages and limits the number of samples to those within a single passage, thus reducing the statistical power. Given the breadth of this study, encompassing nearly 10,000 proteins, after correcting for multiple comparisons, the results indicate that very few, if any, proteins are differentially abundant in each passage (**Figs. S7, S8**). Although various types of analysis of variance tests could be explored for comparisons across more than two groups, a more flexible and suitable choice is a linear mixed-effects (LME) model.

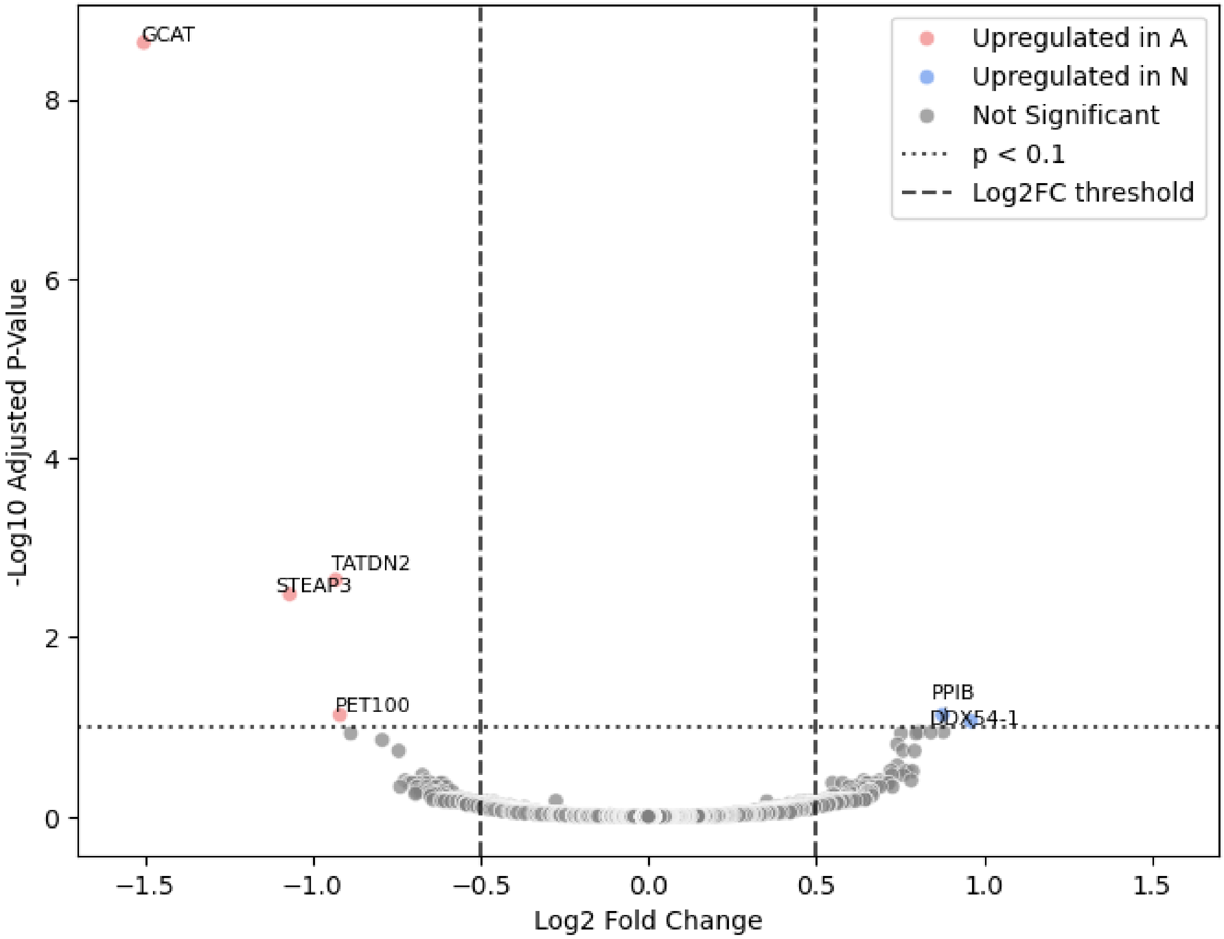


**Figure S6.** Volcano plot of all passages of A *versus* N using a two-sided t-test. All p-values shown are adjusted using Benjamini-Hochberg multiple tests correction.

**N** ***vs.*** **A**

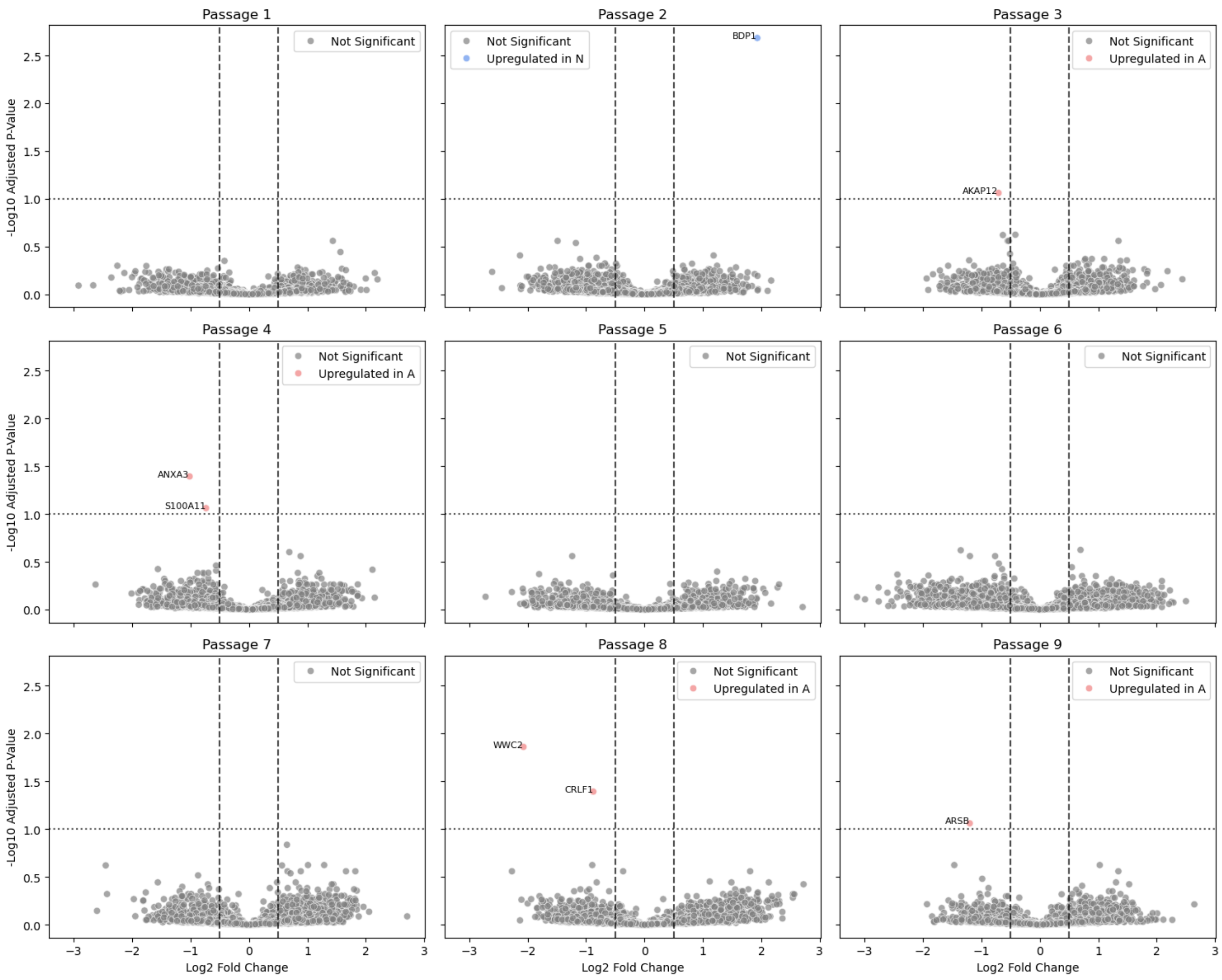


**Figure S7.** Volcano plots of passages 1-9 for comparing conditions N *versus* A. All p-values are adjusted using Benjamini-Hochberg multiple tests correction.

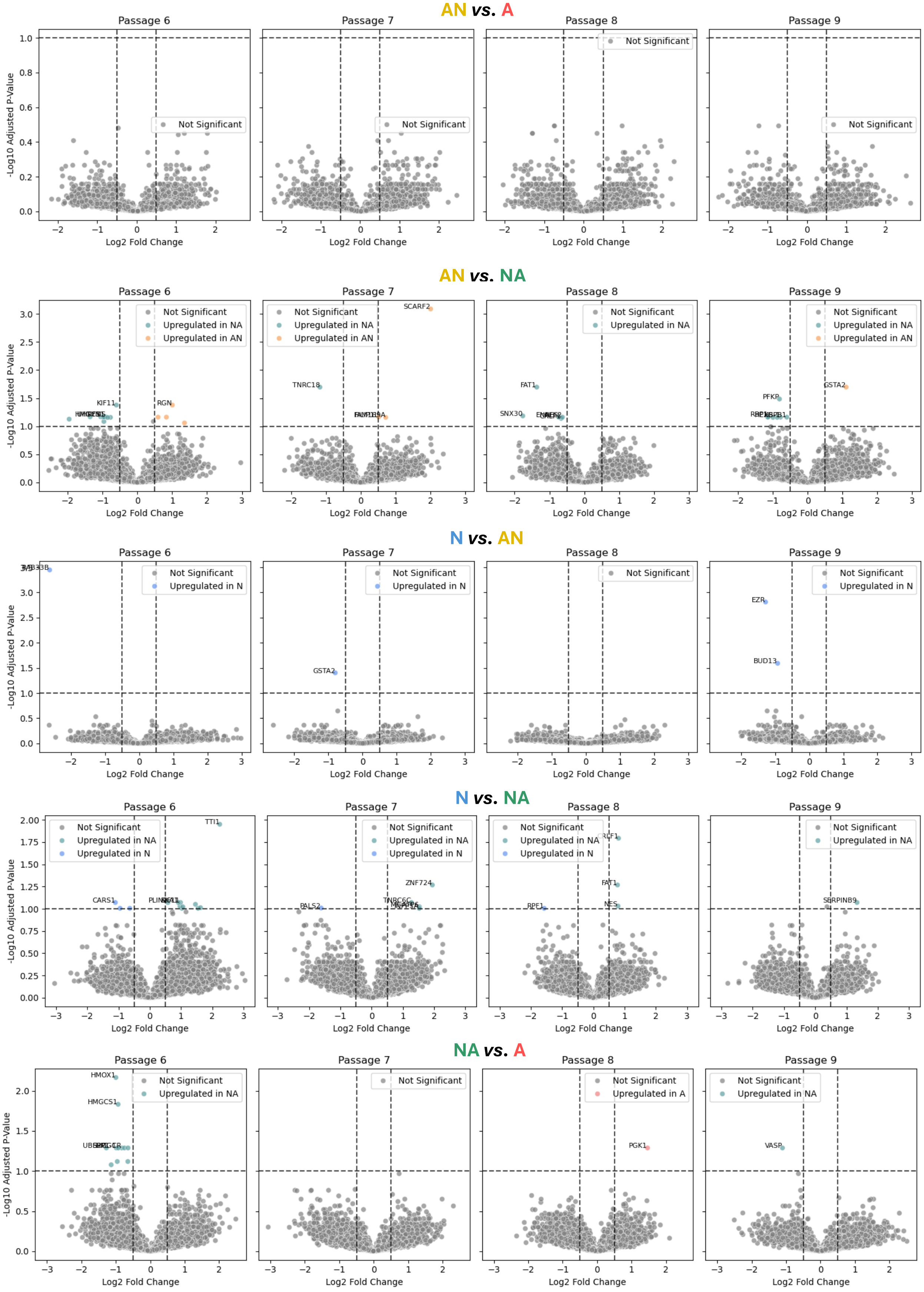


**Figure S8.** Volcano plots of passages 6-9 comparing crossover conditions. All p-values are adjusted using Benjamini-Hochberg multiple tests correction.

**Table S1.** The number of significantly differentially abundant proteins found in each passage/condition shown in Figs. S7,S8.

| Comparison | Antibiotic Treatment | Differentially abundant proteins | | | | | | | | | |
|---|---|---|---|---|---|---|---|---|---|---|---|
| | | **P1** | **P2** | **P3** | **P4** | **P5** | **P6** | **P7** | **P8** | **P9** | **Total (by condition)** |
| N *v* AN | Stop | - | - | - | - | - | 1 | 1 | 0 | 2 | **4** |
| A *v* N | Stop | 0 | 1 | 1 | 2 | 0 | 0 | 0 | 2 | 1 | **7** |
| A *v* AN | Stop | - | - | - | - | - | 0 | 0 | 0 | 0 | **0** |
| NA *v* AN | Either | - | - | - | - | - | 16 | 4 | 5 | 8 | **33** |
| NA *v* A | Start | - | - | - | - | - | 11 | 0 | 1 | 1 | **13** |
| N *v* NA | Start | - | - | - | - | - | 13 | 5 | 4 | 1 | **23** |
| Total (by passage) | | 0 | 1 | 1 | 2 | 0 | 41 | 10 | 12 | 13 | **80** |

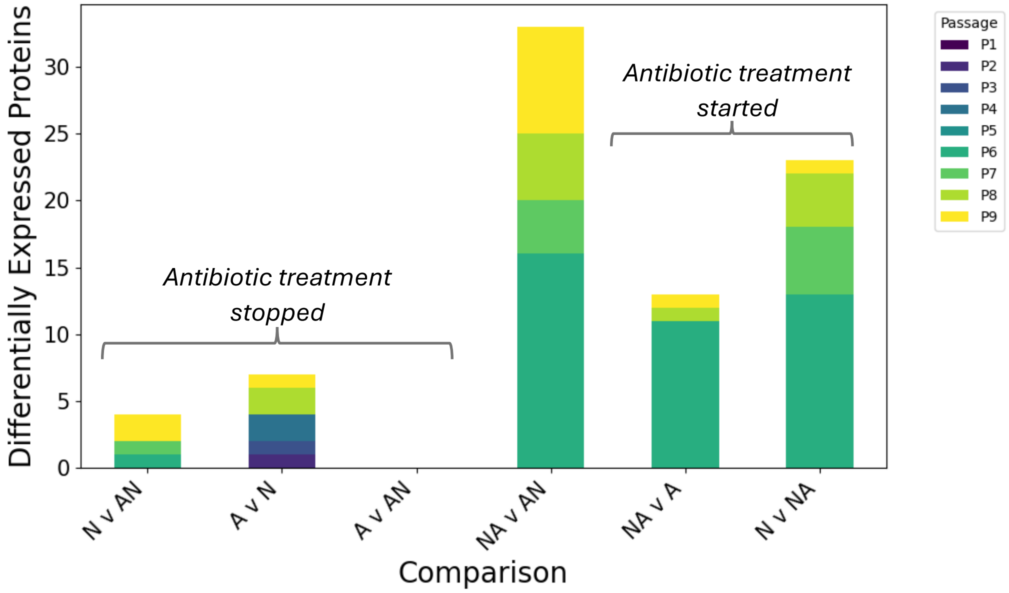


**Figure S9.** The number of differentially expressed proteins by condition and cross-over treatment, stacked by passage.

**Supplementary Note 3**

The choice of whether to include a variable as a fixed or random effect depends on the experimental context.[3] Here, we are interested in the consistent, or average, effects of antibiotic treatment conditions on protein abundance, regardless of a specific passage. Thus, the condition should be considered a fixed effect in our context, while the passage should be regarded as a random effect.

For LME models, the random effects can be specified in the model as random intercepts, random slopes, or a combination thereof. Because we have collected data across multiple passages, a reasonable assumption is that each passage has slight variations in baseline protein quantities simply due to minor differences in the manner the cells were handled, seeded, and grown in each passage. This could include loss of cells and thus differences in proliferation and confluency unrelated to antibiotic treatment, as noted for samples in passages five and six. For example, while we expect all five replicates of passage $A_1$ to have similar proteomic profiles, the proteomes of samples from passage $A_1$ *versus* $A_2$ may vary randomly. To allow for different passages to have different baseline proteomes, a random intercept model is appropriate. Random slopes can also be used to allow for different magnitudes in responses. For example, a protein may respond differently to each condition depending on the passage number it is in. A random slope allows for such variation. By accounting for random effects, the fixed effects of the model are more representative of the average effect of condition on proteomes, *i.e.,* what effects would be present regardless of if we chose these specific passages. However, we were unsure whether to use a random-intercept or random-slope model.

A common strategy is to start with the simplest linear models and build in additional effects until the model stops improving a desired metric.[3] We thus started with ordinary least squares (OLS) models (*i.e.,* no random effects) and progressed in complexity to linear mixed-effect models. For such models, log-likelihood and the Akaike information criterion (AIC) are two commonly used metrics for model selection.[4] While both metrics measure model fit, the AIC penalizes model complexity. Higher (less negative) values of log-likelihood and lower values of AIC indicate better fit. These metrics were calculated for each model described below.

A simple but reasonable baseline OLS model is that protein expression is a function of condition, *e.g.,* $Protein \sim Condition; Y = \beta_0 + \beta_1 C + \varepsilon$. Here, $\beta_0$ represent an intercept and $\beta_1$ represents a slope, and $\varepsilon$ represents error. Building in complexity, passage can be included as an additional fixed effect, i.e., $Protein \sim Condition + Passage; Y = \beta_0 + \beta_1 C + \beta_2 P + \varepsilon$. However, this assumes that passage has a consistent, average affect across all samples, which is not realistic or interesting given the experimental context. Lastly, we can instead consider an interaction effect with condition, or $Protein \sim Condition \cdot Passage; Y = \beta_0 + \beta_1 C + \beta_2 P + \beta_3 (CP) + \varepsilon$. This allows for passage and condition to have an interaction effect. We denote these OLS models as condition-only (C), condition+passage (C_P), and condition*passage (CxP), respectively.

Moving from OLS to LME models allows for the inclusion of random effects. The simplest LME model is $Protein \sim (1|Passage)$, known as a random-intercept only model. This type of model is often used as a baseline comparison for more complex LME models. Because passage differences contributed a large portion of variation as opposed to condition and this model allows each passage to have a different baseline protein abundance, it is expected that this model would perform well as opposed to any OLS model, which tries to find a universal baseline (intercept). Indeed, the RI model exhibited more optimal log-likelihoods and AIC values than any OLS model (**Fig. 3b,** main text).

Adding a random slope slightly worsened model performance. This result was somewhat counterintuitive, as we may expect that proteins across passages would respond to antibiotics differently, *i.e.,* the response to the first time a protein is introduced to antibiotic-free media ($N_1$) would probably be different than passaging it several more times in the same media ($N_{2\text{-}9}$). However, this was not the case. This logic can be rectified by considering that only passages 1 and 6 likely have significantly different responses, as these are the first passages that contain entirely new conditions. Further, the variation may be accounted for by the random intercept, as the random slopes have similar profiles to that of the random intercepts, but at a lower magnitude (**Fig. S10**). Passages 2-5 and 7-9 likely have very similar responses per protein, as they are passaged under the same conditions. Thus, allowing for random slopes overcomplicates the model with little gain in model performance, as shown by comparable log-likelihoods but a worse AIC for random-slope versus random-intercept-only models.

However, neither of the RI or RI_RS models include condition. Next, we can add condition as a fixed effect (FC) to each of the latter two models. This informs the model of our full experimental design; we are looking for consistent (*i.e.,* fixed) variations in proteins across treatment groups, while allowing for different fluctuations in passages to be accounted for simply as noise. Such fluctuations due to passage are not of interest in this study. For example, suppose a protein varies in a single passage. In that case, while this may be interesting for deciphering drug response mechanisms, we are primarily interested in scenarios where proteomes are chronically altered due to antibiotic use. Thus, we want to see changes that are present regardless of the effect of a single passage. This is also why studies that only consider a single passage difference are less experimentally relevant. In general, researchers either 1) always use antibiotics, 2) never use antibiotics, or 3) in the latter case, treat cultures acutely with antibiotics to counteract bacterial contamination before reconditioning them to non-antibiotic conditions. Examining only one differential passage is an unlikely scenario that is rarely encountered in cell culture. The more relevant results are 1) what proteins are chronically different across conditions for multiple passages, and 2) how quickly, if at all, do transient changes rectify themselves (*i.e.,* how long should cell cultures in varying antibiotic supplements be reconditioned before they can be compared with stable proteomes?). A similar trend is observed for fixed condition LMEs with either random intercept, or random intercept and slope, in that the latter performs worse. The similar explanation as above remains. Further, the fixed condition improves log-likelihood over both cases, but AIC remains relatively consistent. Regardless, the results of differential abundance for FC_RI_RS were still largely the same as FC_RI (**Figs. S11, S12**).

Taken together, we found that in general, while the OLS models improved with model complexity (*i.e.,* informing the model of passage), any LME effect model out-performed all OLS models. Within the LME models, the RI and FC_RI models had the most optimal AIC scores. However, the log-likelihood was slightly highest for the FC_RI model. Given the context and variables of interest in the experiment, the FC_RI model provided the best trade-off of suitability and complexity, so we chose this model (**Fig. 3b,** main text) for all remaining analyses unless otherwise noted.

We then plotted the distribution of the random effect coefficients (*i.e.,* random intercepts; **Fig. 3c,** main text). Before swapping conditions (passages 1-5), the variation decreases and is stable by approximately passage 3. Between passages 5 and 6, where the conditions were swapped, a similar trend is noted; by roughly the third passage post-swap (passage 8), the variation is relatively constant, as seen in subsequent passage 9, compared to the following passages. This suggests that, when controlling for the effects of treatment group and passage, the changes induced by antibiotic treatment *versus* non-antibiotic treatment are stable after ~3 passages in a constant condition.  This trend was consistent across random intercept models (**Fig. S10a-c**). A separate study in B16/F10 melanoma cells reported that changes in melanogenic activity in PenStrep-treated cells returned to baseline within as little as 10 hours after reconditioning in non-antibiotic-containing media.[5] Thus, it may be recommended to grow HepG2 cells for at least three passages when switching media conditions to avoid confounding effects.

The distribution of the fixed effect coefficients is shown (**Fig. 3d,** main text). In all fixed effect models, A was used as the reference group and thus represents the intercept or baseline values. The remaining conditions are relative to this baseline. The largest effects are generally in NA and AN, the crossover conditions. In general, condition NA had the largest variation in fixed effects, while condition A had the lowest. This trend makes intuitive sense, as condition A was the baseline condition; cells were in this condition prior to the start of the experiment. The fact that NA shows the great variation in fixed effects is supported by **Fig. S9** and **Table S1**. When comparing *t*-test results pairwise for each condition and passage, we found that starting antibiotic treatment produced more differentially expressed proteins than does stopping antibiotic treatment.

We note that some proteins were consistently identified by various *t*-tests and LME models as differentially expressed (*e.g.,* GCAT) (**Fig. S6, S11**). We also investigated the alternative model which included random slope, FC_RI_RS. Although more proteins were differentially expressed in this model, they were still related to endoplasmic reticulum (ER) stress, chaperone proteins, and various metabolic changes, as shown by the consistent pathway analysis (**Fig. S12**). These results bolster confidence are results are not due simply to the choice of modeling parameters.

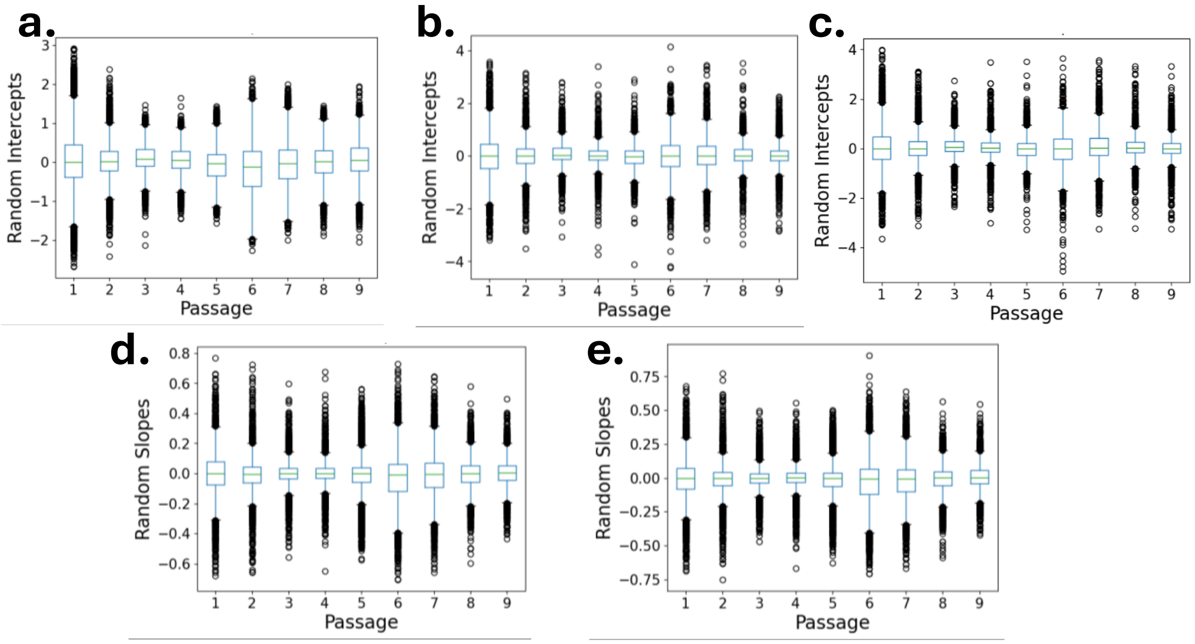


**Figure S10.** Random and effect plots for alternative linear mixed effect models (LME) using **a)** random intercepts only (RI), **b)** random intercepts and random slopes (RI_RS), and **c)** fixed condition with random intercepts and random slopes (FC_RI_RS). The corresponding random slope distributions are shown for **d)** RI_RS and **e)** FC_RI_RS.

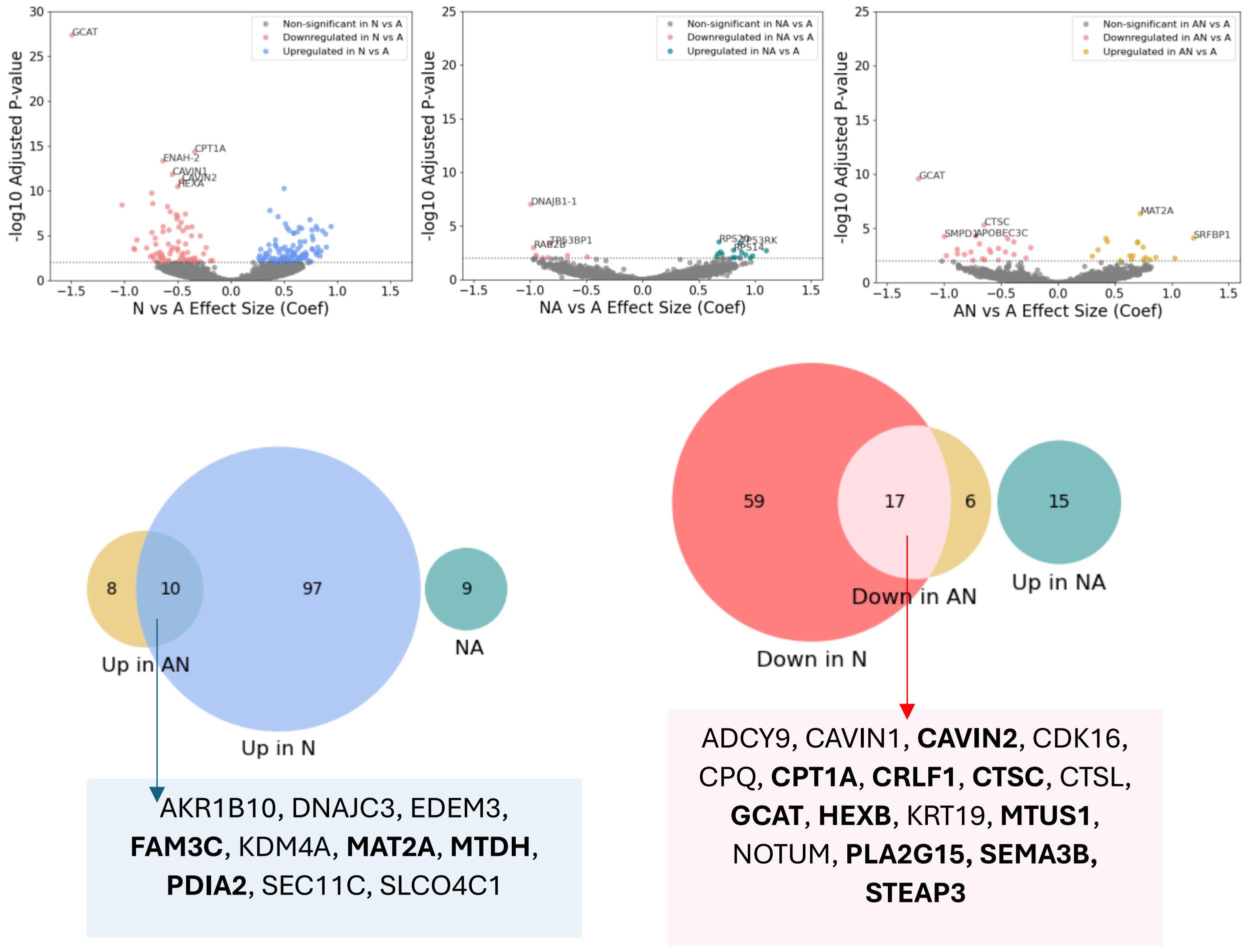


**Figure S11.** Recreated Fig. 3 (main text) using FC_RI_RS results. Bolded proteins represent exact matches between the FC_RI and FC_RI_RS models.

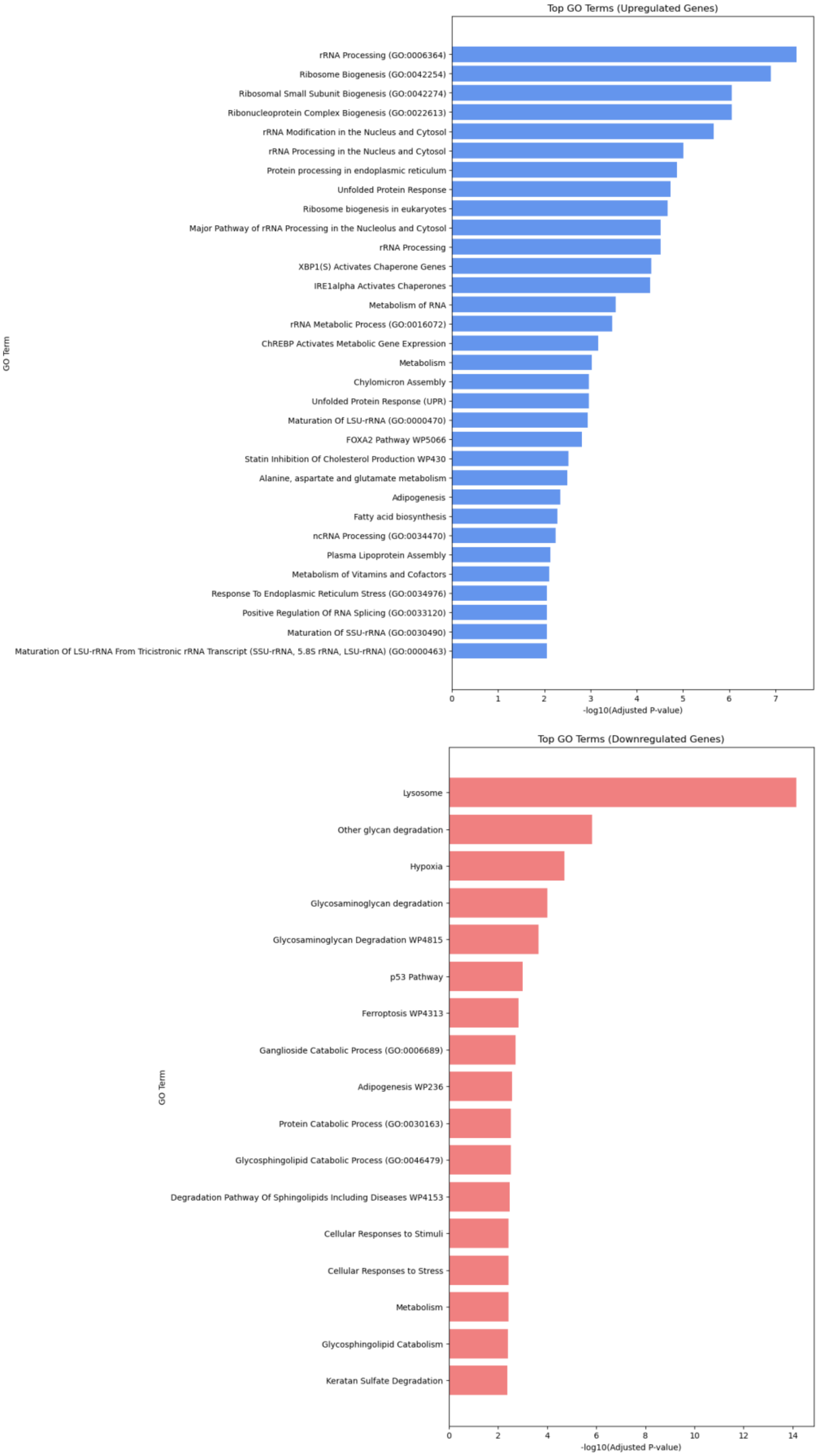


**Figure S12.** Recreated Fig. 4 (main text) using FC_RI_RS results.

**Table S2.** Summary of relevant studies on effects of antibiotics treatment in mammalian cell culture. AmB = amphotericin B, DE = differentially expressed/abundant.

| Study | Technique | Antibiotic(s) | Cell line(s) | Passages/Exposure | Total DE genes/ proteins | p-value Cutoff | Total detected genes/proteins |
|---|---|---|---|---|---|---|---|
| *Current work* | LC-MS/MS | PenStrep | HepG2 | 9 passages (29 days) | 383 (139) | < 0.1 (<0.01) | >10,000 |
| *Ryu et al.* | RNA-seq, ChIP-seq | PenStrep | HepG2 | 2 passages (21 days) | 209 | < 0.1 | >10,000 |
| *Llobet et al.* | Colorimetric/ fluorescence assays | PenStrep, AmB, Gentamicin | Human adipose derived stem cells | 21 days | n/a | < 0.05 | n/a |
| *Mathieson et al.* | 2D-DIGE, MALDI-MS/MS, cell cycle analysis | PenStrep, AmB, Gentamicin | MCF7 | 8 passages (split at $5^{th}$ passage, from anti-anti) | Up to 91 | n/a | 488 |
| *Elliot et al.* | RT-qPCR, colorimetric/ fluorescence assays | Gentamicin | MCF7, MCF12A, MDA-MB-231 | Up to 7 days | 6 | < 0.05 | 8 |
| *Han et al.* | RNA-seq | PenStrep, Gentamicin | Mouse blastocysts | Up to 96 hr | 1,800 | < 0.05 | >10,000 |
| *Relier et al.* | Colorimetric/ fluorescence assays | PenStrep | Six cancer cell lines | Up to 2 weeks | n/a | n/a | n/a |
| *Kalghatgi et al.* | qPCR, colorimetric/ fluorescence assays | PenStrep, ciprofloxacin, ampicillin, kanamycin, tetracycline, spectinomycin | MFC10A and wild type mice | Up to 96 hr | 5 | < 0.05 | 5 |

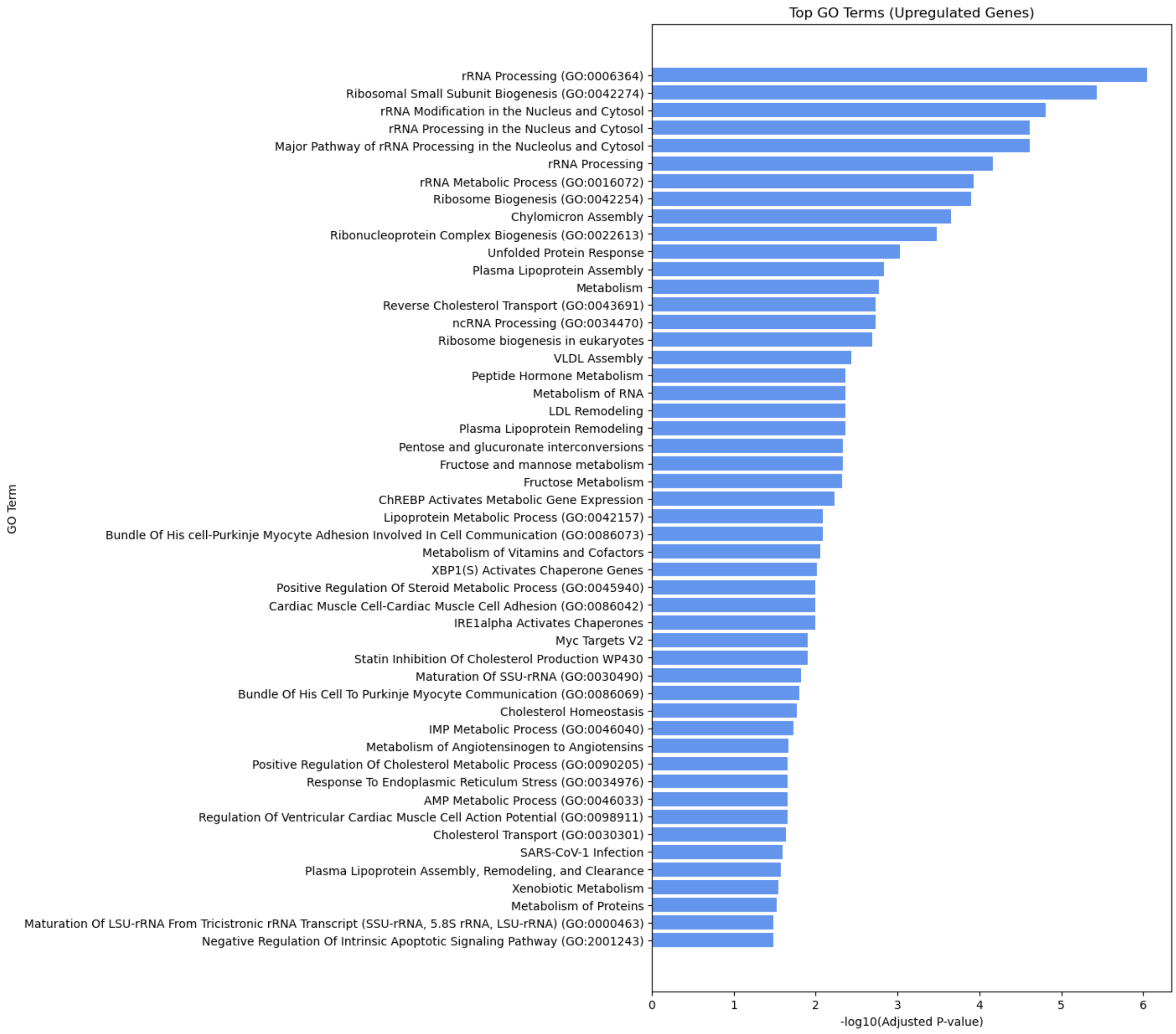


**Figure S13.** Unfiltered GO term enrichment results, upregulated in N.

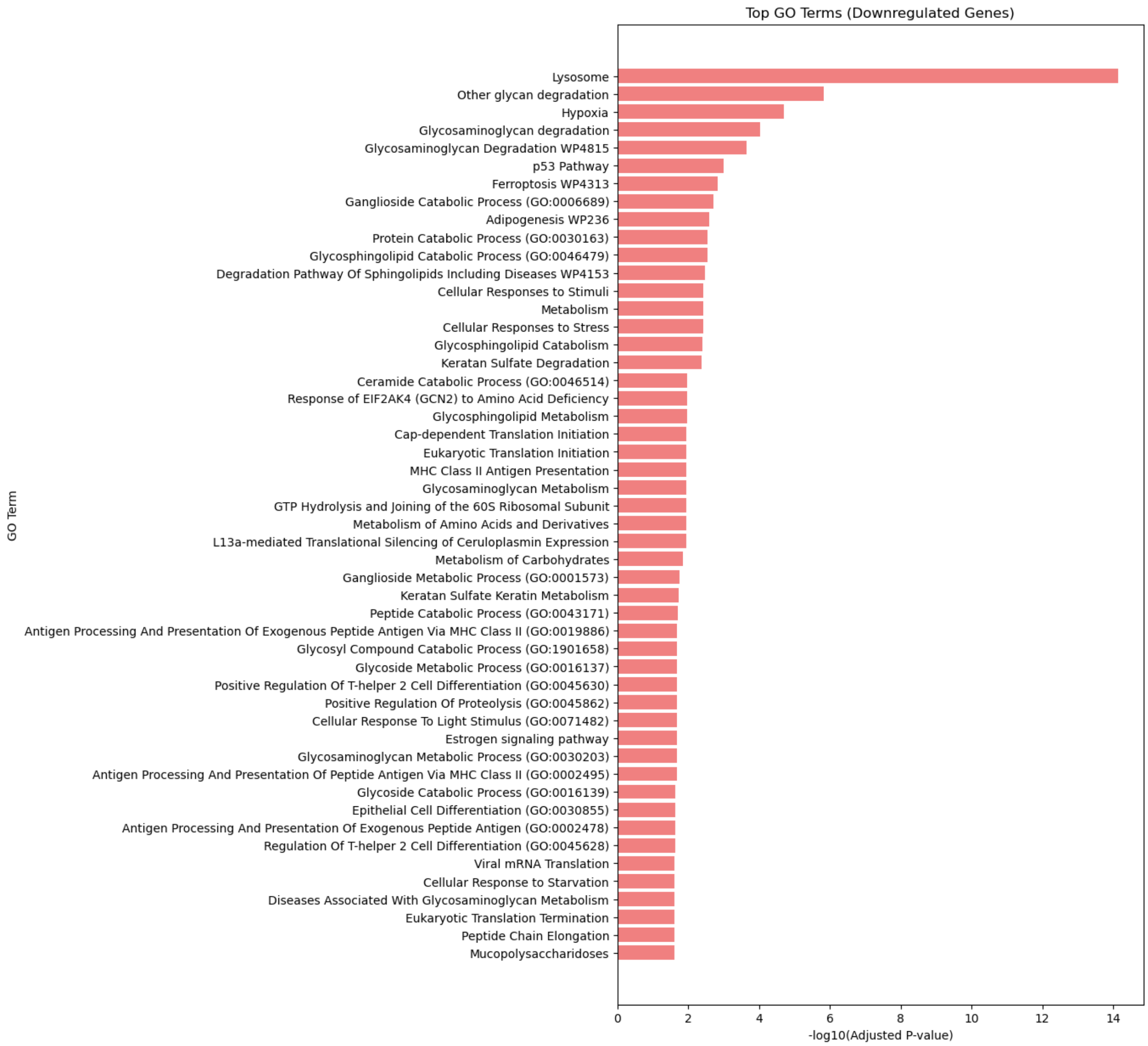


**Figure S14.** Unfiltered GO term enrichment results, upregulated in A.

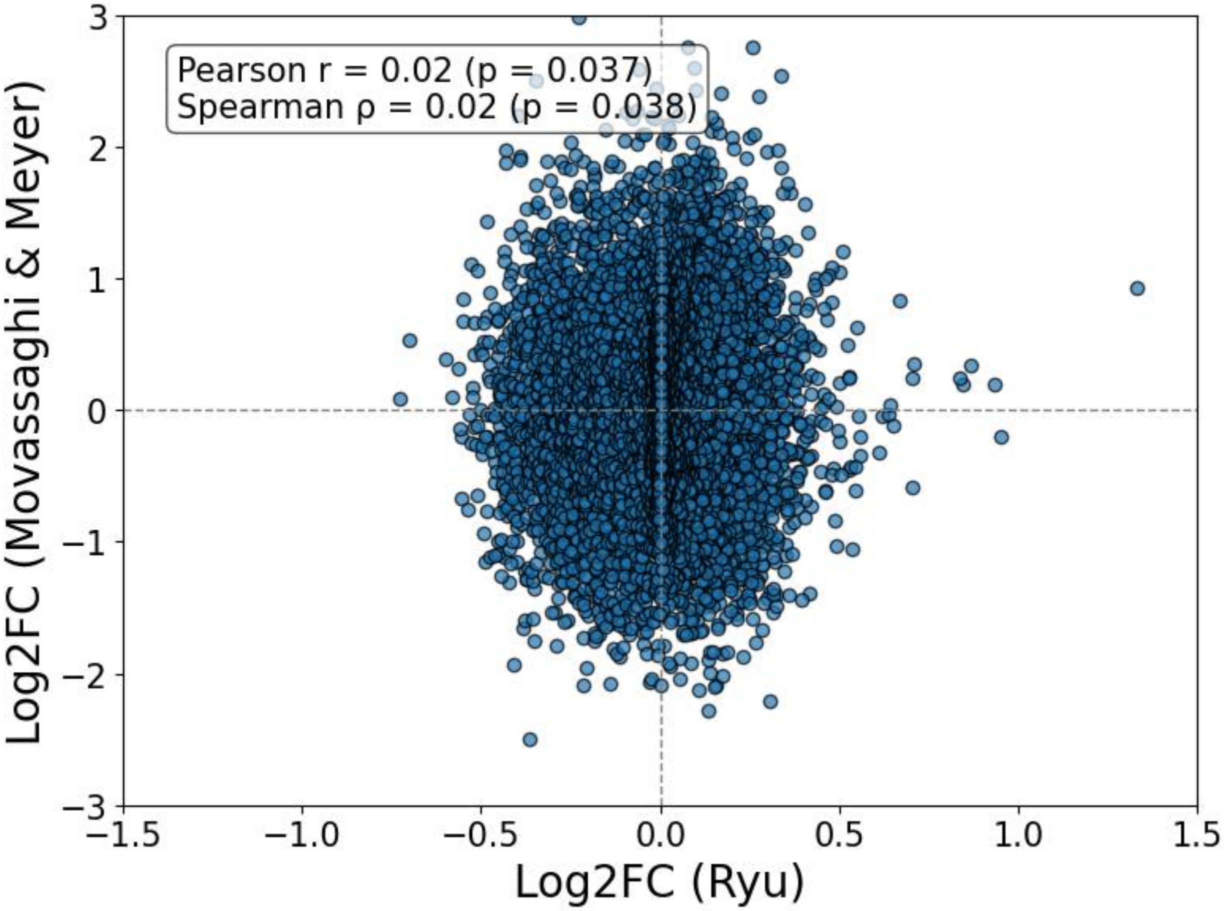


**Figure S15.** Comparison of passage 6 of A *versus* N log2 fold changes with those reported by Ryu *et al.*
